# Exploring molecular signatures of senescence with markeR, an R toolkit for evaluating gene sets as phenotypic markers

**DOI:** 10.64898/2025.12.05.692517

**Authors:** Rita Martins-Silva, Alexandre Kaizeler, Nuno L. Barbosa-Morais

## Abstract

Many biological processes, including cellular senescence, manifest as diverse phenotypes across cell types and conditions. Lacking definitive markers, researchers often rely on the expression of sets of genes to identify these complex states. However, multiple approaches exist to summarise gene set expression into quantitative metrics (*i.e.*, signatures), each with distinct strengths and limitations, and we know of no consensual framework to systematically evaluate their performance across datasets. We therefore developed markeR, an open-source, modular R package that evaluates gene sets as phenotypic markers using scoring and enrichment-based approaches. markeR generates interpretable metrics and intuitive visualisations for benchmarking gene signatures and exploring their associations with study variables. As a case study, we applied markeR to 9 published senescence-related gene sets across 25 RNA-seq datasets, 6 human cell types and 12 senescence-inducing conditions. Gene set performance varied widely: some signatures (*e.g.*, SenMayo) were robust senescence markers across contexts, while others (*e.g.*, MSigDB sets) performed poorly. We further applied markeR to 49 GTEx tissues, revealing tissue- and age-related differences in senescence-associated signals. Together, these findings emphasise the difficulty of characterising molecular phenotypes and demonstrate markeR’s potential for the systematic evaluation of gene sets in various biological contexts.

**GRAPHICAL ABSTRACT:** 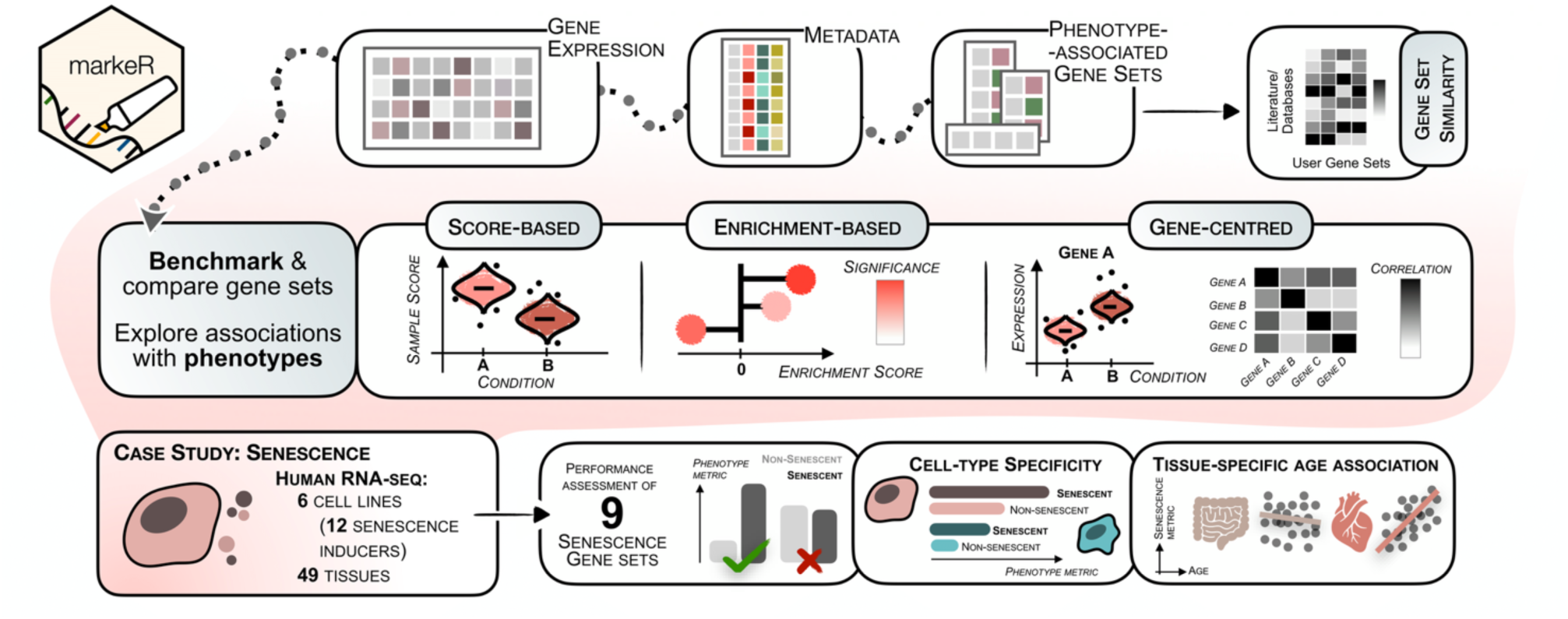

## INTRODUCTION

Originally thought to be a by-product of ageing, cellular senescence is now recognised as a tightly regulated biological programme that plays critical roles from embryonic formation to tissue repair and tumour suppression throughout the life of almost all organisms (1–3).

Described as a stable and irreversible cell cycle arrest under physiological conditions, it is an intrinsic stress response of the cell induced by various factors, including telomere shortening, oncogene activation and DNA damage (4–6). This cell cycle arrest is most often accompanied by profound changes in metabolic activity and secretory phenotype of the cell, the senescence-associated secretory phenotype (SASP), composed of cytokines, growth factors and proteases, all contributing to multiple roles, from immune recruitment to induction of senescence in neighbouring cells (7).

At the molecular level, senescence is highly context-dependent and is regulated by different players such as p53, p16^(INK4A)^ and p21^(CIP1/WAF1)^, which in turn drive distinct gene expression programmes (8,9). This diversity, coupled with the ability of different types of stress, such as DNA damage or oncogene activation, to induce senescence in a cell-type-specific manner, results in a heterogeneous and complex blend of phenotypes rather than a single, well-defined state (10). Such complexity makes it challenging to identify a universal marker, requiring the use of gene sets, rather than individual genes, as molecular signatures of senescence (9,11). This challenge is not unique to senescence, as many biological processes also present as diverse cellular states rather than discrete phenotypes.

Several gene sets have been proposed for senescence, typically derived from transcriptome analyses of senescent cells, or simple curation of genes already reported in the literature as associated with senescence (4,12–15). These gene sets often perform well as senescence markers in the specific context in which they were developed, but lack broader application otherwise. Furthermore, there is no consensual and robust strategy for combining such genes in a meaningful and reliable metric (*i.e.*, a gene signature) for the biological state. Common approaches are based on expression averages (*e.g.*, simple log-median (16), DAS and MSS (17)), rank reflecting relative rather than absolute expression (*e.g.*, GSEA (18), ssGSEA (19), CS score (20), SRRS (21)), or weight-based scoring (*e.g.*, CSS (22)).

Some studies have proposed universal transcriptomic senescence scores (23), but they often lack controls, such as samples of quiescent cells, to distinguish senescence from other non-proliferative states, and extrapolate their findings to complex biological contexts, such as human tissues, where direct validation of the senescent phenotype remains challenging.

While benchmarking studies have compared signature-scoring methods in specific contexts (24), each has its own strengths and limitations, and, to our knowledge, there is no flexible platform allowing researchers to systematically test, compare and apply multiple approaches with their gene sets of interest.

Herein, we present markeR, an R package that provides methods to quantify how well a gene set marks a phenotype by transforming its genes’ expression profiles into various metrics and assessing their robustness. markeR can be used for (1) comparing gene sets for consistency and specificity in marking a phenotype (benchmarking), and (2) quantifying the gene set’s expression and exploring its relationship with phenotypic variables (discovery). To demonstrate its applicability, we curated a compendium of publicly available gene expression datasets and senescence-related gene sets. While senescence serves as a case study due to its complexity and relevance, the framework is broadly applicable to other contexts. markeR is freely available in Bioconductor at https://bioconductor.org/packages/markeR.

## MATERIAL AND METHODS

`markeR` (v1.0.0) was developed in `R` (v4.4.3) with a modular design to support easy integration of new features and functionalities (Figure 1). It was designed for users looking for both a quantitative framework with different methods to characterise a given phenotype, and a fully customisable visualisation component to suit their needs. It requires as input:

1. A list of gene sets associated with the phenotype of interest. For each gene set, information on the expected direction of regulation change (*i.e.*, up- or down-regulation) of each gene upon phenotype may or may not be included;
2. A pre-processed (*i.e.*, filtered and normalised, but not log-transformed) gene expression matrix;
3. A sample metadata table.

**Figure 1.**
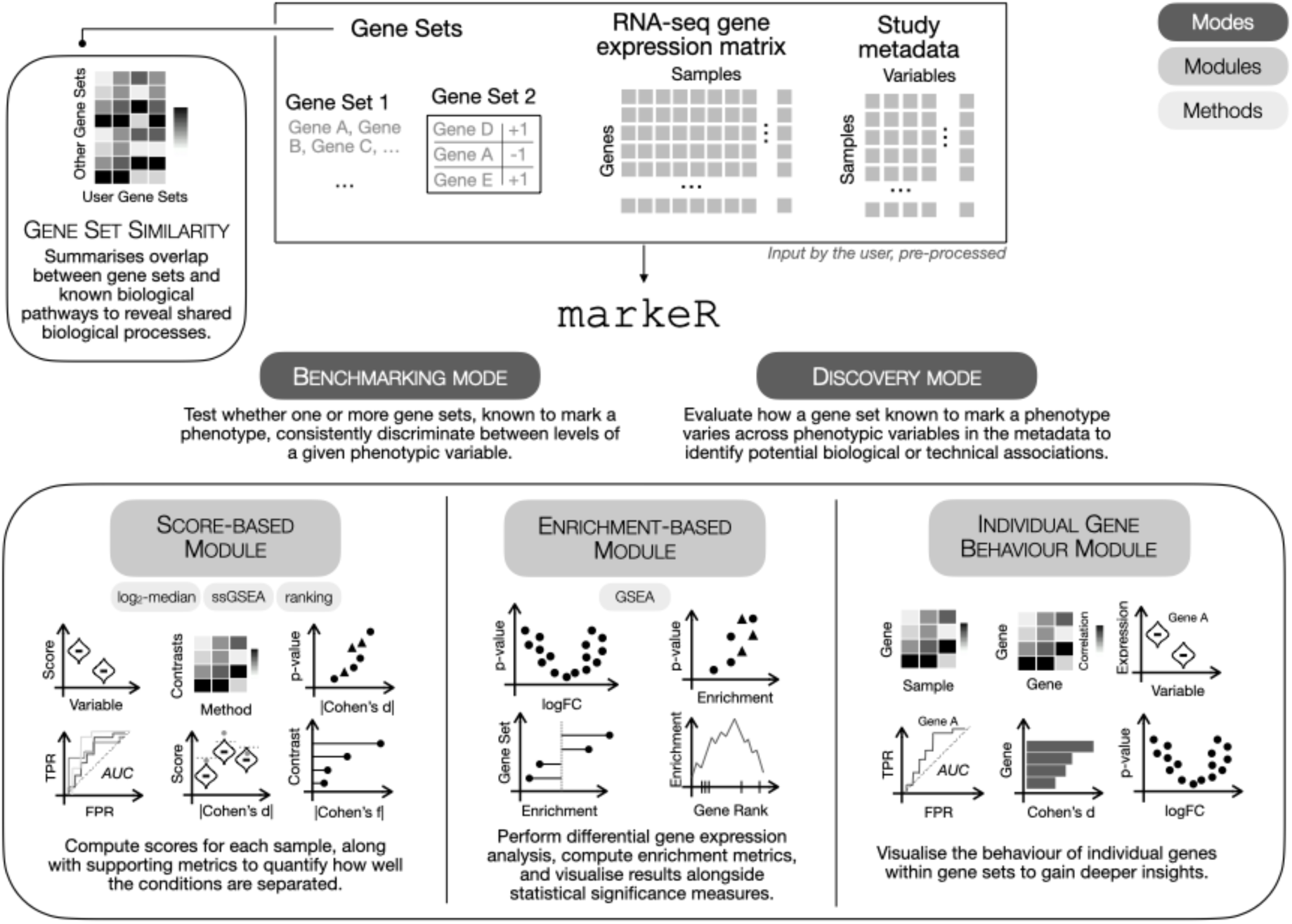
**– Overview of markeR**. markeR requires three main inputs: (1) one or more gene sets putatively associated with a phenotype of interest, to be provided with or without directionality (e.g., gene set 1 is made of genes with no information on the expected direction of regulation, therefore assumed to be up-regulated upon phenotype; gene set 2 includes genes explicitly annotated as up- or down-regulated upon phenotype); (2) a pre-processed (filtered for low expression genes and normalised) RNA-seq gene expression matrix; and (3) sample-level metadata. Users can choose between two analysis modes: benchmarking, to test the robustness of signatures in discriminating between phenotypic groups of samples, and discovery, to profile associations between a given signature and selected (from metadata) phenotypic variables. Regardless of the mode, the quantitative evaluation of signatures as phenotypic markers can be done using score-based or enrichment-based methods. The contribution of individual genes to the respective gene sets’ performance can also be explored. markeR also quantitatively compares between user-defined and curated reference gene sets, to enhance biological interpretability.

### Modules for gene set-based phenotype characterisation

#### Score-based approaches

- *Log_2_-median-centred score.* The **log_2_-median** method first applies a log₂ transformation to the expression values, followed by median-centring each gene’s expression across all samples. Each sample’s score is the mean of the median-centred expression values therein of the genes in the set (example of its use in (16)). Each gene’s contribution is evaluated relative to its typical expression, and the resulting score reflects the coordinated expression of genes within the gene set. For bidirectional gene sets, i.e., that include information on the expected direction of change of each of its genes upon phenotype, the sample score is the mean of median-centred expression for the upregulated subset of genes minus that of the downregulated subset.

- *Single sample Gene Set Enrichment Analysis (ssGSEA) score.* Single sample Gene Set Enrichment Analysis (ssGSEA) was implemented using a modified version of the `GSVA` package’s `gsva` function (19), based on the original Gene Set Enrichment Analysis (GSEA) method (18). ssGSEA ranks all genes by their expression within each sample and computes a running-sum statistic over the ranked list. For unidirectional gene sets (i.e., with no information on the expected direction of each gene’s regulation upon phenotype), the ssGSEA sample score reflects overall coordinated expression changes of the genes in the set. For bidirectional gene sets, the score is calculated as the difference in ssGSEA scores between the upregulated and the downregulated subsets of genes.

- *Ranking score.* The ranking method uses a non-parametric approach based on the relative expression of genes within a set. For each sample, all genes are ranked by expression. The score is the sum of the ranks of the genes in the gene set, divided by the number of genes in the set. The individual ranks of putatively downregulated genes in bidirectional gene sets are multiplied by −1.

*Enrichment-based approach.* markeR uses GSEA in its enrichment-based module (18). Linear-model-based differential gene expression analysis for the phenotypes of interest is performed with limma (25) (v3.62.2). If the phenotypic variable of interest is categorical, (*e.g.*, treatment, genotype) the design matrix is constructed with model.matrix(∼0 + variable)(model without intercept). Pairwise comparisons between levels of this variable are specified using a contrast matrix. For continuous variables, the model (with intercept) ∼1 + variable is used instead. Alternatively, a custom design matrix modelling the user’s biological questions can be provided. Genes are then ranked according to differential expression statistics:

- for gene sets without specified direction of expression changes upon phenotype (unidirectional), the B-statistic, *i.e.*, the empirical Bayes posterior log-odds of differential expression (26), irrespective of the direction of change;

- for bidirectional gene sets, the empirical Bayes moderated t-statistic (26), *i.e.*, representing the ratio of estimated effect size to its standard error, multiplied by −1 for putatively downregulated genes.

GSEA is then performed using the fgsea package (v1.32.4), and interpreting the resulting normalised enrichment score (NES) for each gene set in each contrast (*i.e.*, phenotype)(18) depends on the differential expression statistic used for gene ranking:

- B-statistic: A positive NES indicates the gene set’s enrichment among the most altered genes; a negative NES, mostly uninformative, indicates the gene set’s depletion among the most altered genes (*i.e.*, enrichment among the least altered genes);

- t-statistic: a positive or a negative NES indicate the gene set’s over- or under-activation, respectively.

Significance is determined using Benjamini-Hochberg’s False Discovery Rate (FDR) correction for multiple testing of all analysed gene sets. An optional additional FDR correction can be applied to account for the number of contrasts tested per gene set, providing a more stringent control of significance (addressing the question *“Are there significantly altered gene sets in any of the contrasts?”* rather than *“Are there significantly altered gene sets in my contrast of interest?”)*.

*Individual Genes.* markeR provides tools to assess the contribution of individual genes to their sets’ score- and enrichment-based results. Visual and analytical outputs include expression heatmaps and violin plots of gene expression; correlation heatmaps to explore relationships between genes in the set; receiver operating characteristic (ROC) curves and respective areas under the curve (AUC) to evaluate gene-level classification performance; effect size (Cohen’s d) of expression differences between sample groups; and principal component analysis (PCA) of gene expression profiles to assess the gene set’s contribution to phenotype-associated variance.

### Benchmarking and discovery modes

markeR provides two modes for evaluating gene sets as phenotypic markers: **benchmarking** and **discovery**.

**Benchmarking** assesses the robustness of gene sets’ performance across multiple analytical methods. It includes violin plots for distributions of gene set scores across categorical phenotypes, while scatter plots are used when the phenotypic variable is numerical. Volcano plots display the statistical significance (-log_10_ p-values from Student’s t-tests) against the effect size of differences in the distributions of sample scores (Cohen’s d) between each pair of phenotyping groups of samples. Complementary heatmaps colour-code the effect size and annotate them with corresponding p-values. Additional features include ROC and AUC calculations (via the pROC package, v1.18.5), as well as estimation of effect size using Cohen’s f for comparisons involving more than two groups or continuous variables (via the effectsize package, v1.0.0). Some analyses (*e.g.* ROC curves, Cohen’s f heatmaps) apply only when the variable of interest is categorical. To evaluate whether a gene set’s observed effect size exceeds what would be expected by chance for sets with the same characteristics, markeR quantifies gene set-level significance as the false positive rate (FPR), also commonly designated as empirical p-value, by comparing the observed effect sizes against null distributions of effect sizes in 100 gene sets, each generated by random selection of the same number of genes and, for bidirectional sets, expected proportion of genes up- or down-regulated upon phenotype. Results are visualised as violin plots of randomly generated Cohen’s d or f values across methods, overlaid with dots for the empirical values and a 95^th^ percentile significance threshold. For enrichment-based analyses, results from linear models are summarised using scatter or lollipop plots of NES values and their statistical significance. plotEnrichment (fgsea package) is used to plot the running enrichment score across the ranked gene list, highlighting the ranks of gene set members. Volcano plots, displaying significance against effect size, further contextualise gene set members within the broader expression landscape and help assess their robustness as markers.

**Discovery** assists in hypothesis generation by profiling associations of scores or enrichment values with selected phenotypic or clinical variables. Score distributions are summarised alongside effect sizes: Cohen’s d for pairwise comparisons or Cohen’s f otherwise. Cohen’s f was calculated for each variable using a linear model of the form score ∼ variable; for continuous variables with more than two levels, significance was assessed using the t-test for the regression slope, whereas for categorical variables, significance was obtained from an ANOVA F-test comparing group means. NES values are summarised in lollipop plots for selected contrasts, with tunable resolution. Dots are colour-coded by FDR.

Benchmarking includes the most comprehensive feature set, allowing variables of interest identified in Discovery to be seamlessly further tested. All results are visualised on plots generated by ggplot2 (v3.5.2) and ComplexHeatmap (v2.22.0). Depending on the function, plots may be combined using ggpubr (v0.6.0), grid (4.4.3) or cowplot (v1.1.3).

#### Similarities with other gene sets

To get insights on the biological relevance of the gene sets, similarity to other sets can be assessed using two metrics: the Jaccard index and the odds ratio (OR). The Jaccard index (JI) is calculated as the size of the intersection divided by the union of two gene sets. The OR is derived from a 2×2 contingency table based on a user-defined gene universe, with statistical significance given by the p-value of the one-sided Fisher’s exact test. Comparisons are made between user-defined and curated reference gene sets, which may include other user-defined and MsigDB collections (18) (accessed via the msigdbr package, v24.1.0), and results can be filtered by minimum Jaccard index or a combination of odds ratio and p-value.

Results are displayed as a heatmap (ggplot2 v3.5.2), with colours representing either OR on a log_10_ scale or the JI. In this study, we focused on the Hallmarks collection and used a gene universe comprising all genes annotated in The Human Protein Atlas. We displayed gene sets with FDR ≤ 0.05 and odds ratio >1.

### Senescence RNA-seq datasets

*Dataset Collection.* For evaluating gene sets as senescence markers, we curated human RNA-seq datasets from the Gene Expression Omnibus (GEO) (27) and ArrayExpress (28), assisted by literature review based on PubMed searches. Curation followed a two-step strategy. We first focused on fibroblast-derived datasets, given their widespread use as a tractable model for studying cellular senescence. Inclusion criteria required senescent or quiescent samples, with a minimum of 10 samples per study, excluding studies using cancer cell lines or tumour-derived material (specific queries in **Supplementary File 1 – GEO, ArrayExpress**). All such datasets available as of 14 February 2024 (GEO) and 29 February 2024 (ArrayExpress) were considered. Validation of senescence (*e.g.*, SA-β-gal staining, cell cycle arrest markers) was manually assessed from primary publications, and studies with insufficient or unclear validation were excluded. 16 datasets, comprising 384 samples, met all criteria (**Supplementary File 1 – Final Datasets, Metadata**). To ensure generalisability across cell types, we then considered all non-fibroblast datasets available as of 12 July 2024, excluding studies using cancer cell lines or tumour-derived material (specific queries in **Supplementary File 2 – GEO, ArrayExpress**). 9 datasets, comprising 161 samples, met all criteria (**Supplementary File 2 – Final Datasets, Metadata**). These comprised five cell types: endothelial (4 datasets, 90 samples), keratinocyte (2 datasets, 36 samples), melanocyte (1 dataset, 12 samples), mesenchymal (1 dataset, 12 samples) and neuronal (1 dataset, 11 samples). All samples (fibroblast and non-fibroblast) were categorised into three cellular states according to the respective primary study: proliferative (272 samples), quiescent (33) and senescent (240).

*Data download and alignment.* Data download and quality control were performed in a conda environment (Python v3.11.5). FASTQ files were obtained from GEO using fasterq-dump (v2.11.0; SRA toolkit v2.11.0; NCBI, https://github.com/ncbi/sra-tools) and manually from ArrayExpress. Read quality was assessed using fastqc^1^ (v0.12.1) and summarised using MultiQC (29) (v1.14). All samples passed quality control and were retained for analysis. Reads were pseudo-aligned to the RefSeq reference human transcriptome (NCBI release 109)(30) using kallisto (31) (v0.44.0).

*Preprocessing.* Genes with low expression across conditions were removed by retaining only those with an average of > 70 reads/sample in at least one condition (quiescent, proliferative, or senescent)(**Supplementary Figure 1A**). Normalisation factors were calculated using calcNormFactors from the edgeR package (32), and read counts were subsequently normalised and log_2_-transformed using voom from the limma package (**Supplementary Figure 1B**).

*Batch Effect Correction.* Batch correction was performed using a modified version of the removeBatchEffect function from the limma package, following a strategy similar to Schneider et al., 2024 (33). Dataset ID (*i.e.*, source study) was treated as a technical batch variable and regressed out of the expression matrix. In contrast, sources of variance reflecting biological differences (*e.g.*, cell type, cellular state) were modelled and kept in the analysed data (**Supplementary Figure 1C**).

### GTEx Human Transcriptomic Data

*Data download.* Pre-processed RNA-seq data from The Genotype-Tissue Expression (GTEx) v8 were obtained from the GitHub repository associated with the Schneider et al., 2024 publication (https://github.com/DiseaseTranscriptomicsLab/voyAGEr) (33).

*Correction for cell-type composition.* To assess the potential confounding effect of age-related shifts in cell type composition on the observed age-senescence associations in GTEx data, cell type enrichment scores were estimated for each tissue using xCell2 (34) (v1.2.3) with the pre-trained BlueprintEncode reference, covering 43 cell types derived from Blueprint and ENCODE bulk RNA-seq datasets. For each tissue, the six cell types with the highest score variance across samples were selected as covariates, prioritising those most likely to co-vary with age. A corrected model including these scores was compared with the original age-only model using the enrichment module of markeR, and Benjamini-Hochberg correction was applied separately per model across all 49 tissues. To allow comparison between models, observed NES values were expressed as percentile ranks relative to a null distribution obtained by permuting the age variable 1000 times per tissue, for the uncorrected model.

### Senescence Gene Sets

Senescence gene sets were selected based on resources publicly available as of 13 March 2024 (**Supplementary File 3 – Senescence Gene Sets**). We included five well-cited gene sets empirically derived from different cell types and senescence-inducing stimuli: SAUL_SEN_MAYO, CSGene (15), CellAge (35), SeneQuest (4) and HernandezSegura (13). Additionally, we incorporated four gene sets from MSigDB (18): GOBP_CELLULAR_SENESCENCE, GOBP_NEGATIVE_REGULATION_OF_CELLULAR_SENESCENCE, GOBP_POSITIVE_REGULATION_OF_CELLULAR_SENESCENCE, and REACTOME_CELLULAR_SENESCENCE.

## RESULTS

*markeR is a flexible R package designed to quantify, compare and evaluate gene expression signatures as markers of cellular phenotypes (****Figure 1****). The package supports two main quantification approaches:*

- **Score-based:** A signature score, summarising the collective expression of a gene set therein, is assigned to each sample.
- **Enrichment-based:** Gene Set Enrichment Analysis (GSEA) is run on genes ranked according to differential expression statistics, and a resulting enrichment score is assigned to each gene set for each phenotype.

### Case Study: Cellular Senescence as a Complex Biological Phenotype

We use senescence as a case study for markeR, as it exemplifies a complex phenotype lacking a universal marker. Although several gene signatures have been proposed to identify senescent cells across diverse contexts, their systematic benchmarking is still missing. To address this, we first applied markeR in benchmarking mode to a curated RNA-seq dataset of senescent, proliferative and quiescent samples of human cells, thereby evaluating the robustness as markers of widely used senescence-associated gene sets across cell types and senescence-inducing stimuli. We then demonstrate markeR‘s discovery mode using RNA-seq data of human tissues from the Genotype-Tissue Expression (GTEx) project, where senescence is not explicitly labelled but is hypothesised to increase with age (18,36).

### Gene Sets and RNA-seq Datasets

For the benchmarking analysis, we selected five senescence-associated gene sets from open-source, well-cited studies and four from MSigDB (see Methods). To focus on gene sets capturing core features of senescence, we excluded gene sets exclusively associated with the SASP, given its known heterogeneity (13). The selected gene sets vary in size, and some include annotations of the expected direction of expression change (*i.e.*, if each gene in the set is putatively up- or downregulated in senescence) (**Figure 2A**). Most gene sets were compiled through similar literature curation approaches, often incorporating the same well-established senescence-associated genes, likely explaining their substantial overlap. In contrast, the HernandezSegura and CellAge gene sets were derived independently from

**Figure 2.**
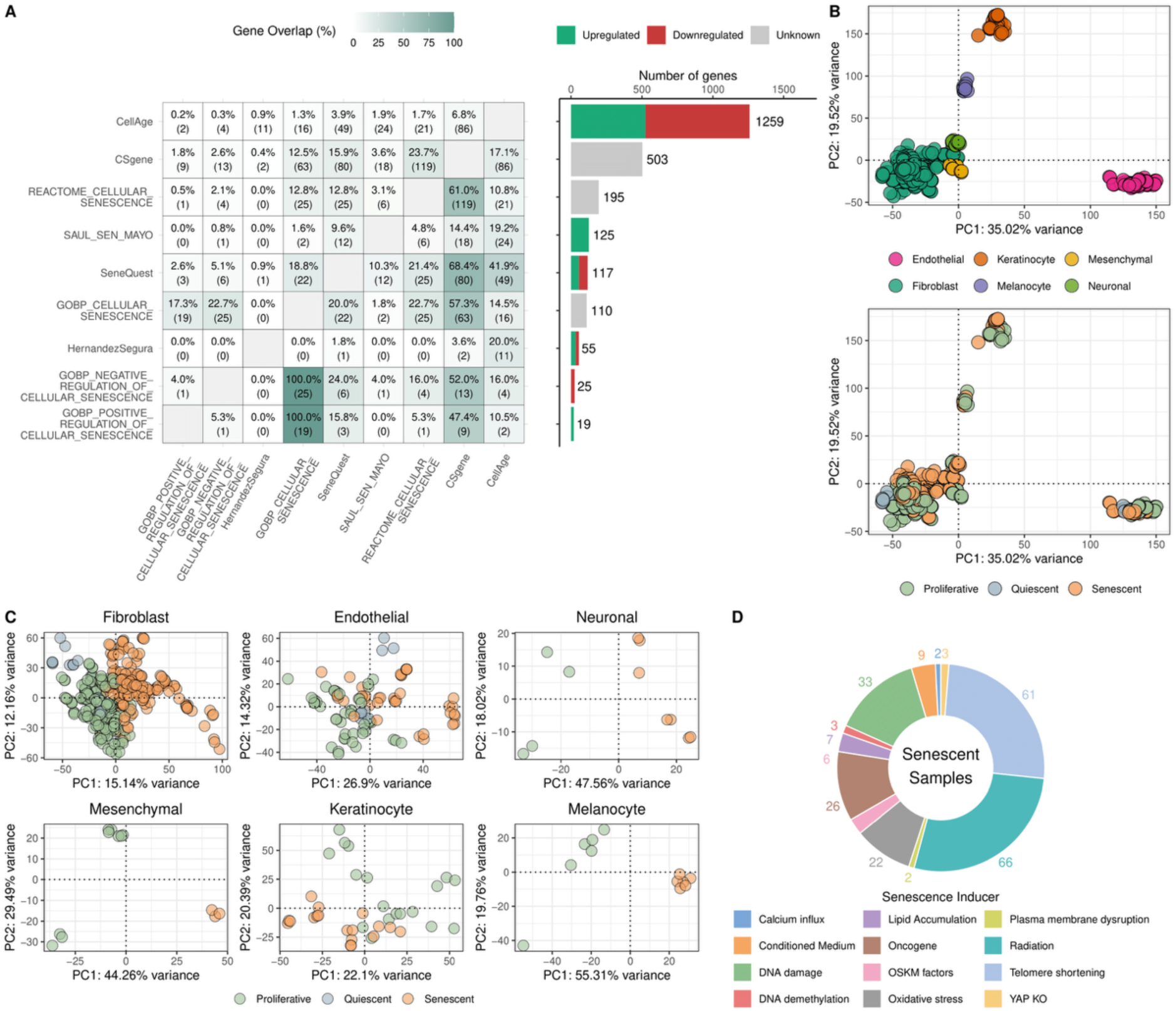
Overview of senescence gene sets and RNA-seq datasets. A collection of 9 publicly available senescence-associated gene sets and 25 RNA-seq datasets, covering 545 samples from six human cell types, was used in this analysis. (A) Heatmap showing the percentage and number (in brackets) of genes of each set in the rows in common with those of sets in the columns (left plot), accompanied by a barplot of the total number of genes per set, coloured by the expected direction of change upon senescence (right plot; upregulated in green, downregulated in red, unknown direction in grey). (B) Principal component analysis (PCA) plots of gene expression showing that, after batch effect correction, most data variance can be attributed to the cell type of origin (top plot) but not cellular state (bottom). Batch correction was performed by regressing out dataset ID as a technical covariate while preserving sources of biological variance (cell type and cellular state); see Methods for details. (C) PCA plots of gene expression per cell type, showing that cellular state is a major source of variance within each cell type. (D) Donut chart of senescent samples per senescence inducer.

RNA-seq and microarray data analyses, respectively, and show minimal overlap with the others (**Figure 2A**). Most sets are enriched for nucleoplasm genes (**Supplementary Figure 2A**), consistent with roles in cell cycle regulation and transcription. Functional analyses (**Supplementary Figure 2B**) reveal significant enrichment in genes involved in proliferation, apoptosis, and immune pathways, aligning with known features of senescence (7).

To evaluate the senescence-marking potential of those gene sets, we curated 23 publicly available human RNA-seq datasets (545 samples) from GEO, ArrayExpress, and manual literature review in PubMed (see Methods), prioritising diversity in cell types and senescence-inducing stressors (**Figure 2B-D**), amongst the main sources of phenotypic heterogeneity in senescence. The overall dataset for analysis includes six cell types (fibroblasts, endothelial, keratinocyte, melanocyte, mesenchymal, and neuronal), twelve distinct stressors, and three cellular states (240 senescent, 272 proliferative, and 33 quiescent samples)(see **Supplementary Files 1** and **2**). Quiescent samples were included as an essential control to distinguish senescence-specific signatures from those associated with general proliferative arrest. After pre-processing all RNA-seq data together (see Methods), cell type was, as expected, the primary driver of gene expression variation (**Figure 2B, Supplementary Figure 1D**), while cellular state (proliferative, senescent, quiescent) emerged as the main source of variance within each cell type (**Figure 2C)**.

### Score-Based Methods Enable Sample-Level Quantitative Marking of Senescence

To quantify the performance of each gene set’s expression in marking senescence phenotypes, we implemented in markeR several score-based methods (log_2_-median, ranking and ssGSEA), each with different assumptions (see Methods) but expected to produce consistent results for robust markers, as observed (**Figure 3A, Supplementary Figure 3**).

**Figure 3.**
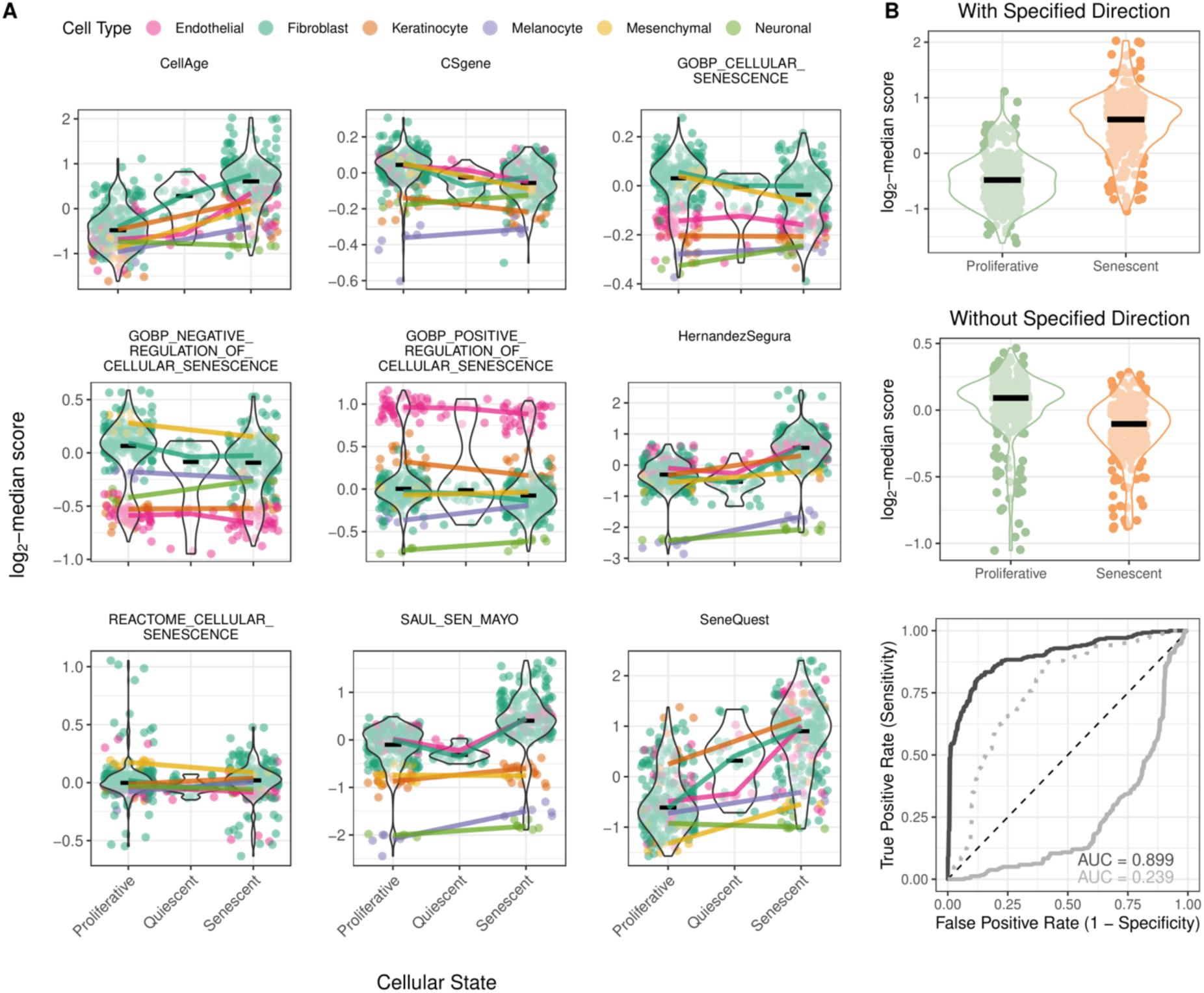
Performance of score-based phenotype quantification across gene signatures. (A) Violin plots of distributions of log_2_-median scores of samples (dots) for each senescence-associated gene signature per cellular state (proliferative, quiescent, senescent), coloured by cell type. Median scores per cellular state for each cell type are connected by coloured lines. (B) Violin plots of distributions of log_2_-median scores of proliferative and senescent samples (green and orange dots, respectively) for the CellAge signature, with (top) and without (middle) the direction of gene expression change upon senescence considered. Bottom panel: Receiver Operating Characteristic (ROC) curves of the log_2_-median score of the CellAge signature to mark senescent in contrast with proliferative samples, with (black) and without (grey; dotted line representing the mirrored ROC) the direction of gene expression change upon senescence considered. The respective areas-under-curve (AUC) are displayed. The loss of directional information can lead not only to a reduction in classification performance but also to misinterpretations.

Some gene sets displayed strong cell type-specific patterns: the scores of CSGene and GOBP_CELLULAR_SENESCENCE increased from proliferative to senescent samples in some cell types but decreased in others, whereas HernandezSegura, SAUL_SEN_MAYO, and CellAge exhibited more consistent trends across cell types. Score distributions varied by cell type, suggesting cell-type-specific baseline expression levels of senescence-associated genes, particularly evident for GOBP_POSITIVE_ and GOBP_NEGATIVE_REGULATION_OF_CELLULAR_SENESCENCE, as observed when samples are stratified by senescence-inducing stimulus (**Supplementary Figure 4**). Stimulus-specific sensitivity in senescence marking was also observed: HernandezSegura and SAUL_SEN_MAYO effectively distinguished senescent from non-senescent samples across stimuli, with particularly strong discrimination in oncogene-induced senescence, while CellAge was more specific to radiation and telomere shortening (**Supplementary Figure 4**). Conversely, REACTOME_CELLULAR_SENESCENCE and GOBP_POSITIVE_REGULATION_OF_CELLULAR_SENESCENCE, showed poor discrimination across conditions. Others, such as SeneQuest, consistently discriminated senescent from non-senescent samples regardless of the inducing stimulus.

HernandezSegura, CellAge and SAUL_SEN_MAYO consistently separated senescence from quiescence, while, CSGene, GOBP_CELLULAR_SENESCENCE and GOBP_NEGATIVE_REGULATION_OF_CELLULAR_SENESCENCE, failed to do so (**Figure 3A**), importantly suggesting they reflect general non-proliferative states rather than senescence-specific programs.

Overall, HernandezSegura, CellAge, SeneQuest and SAUL_SEN_MAYO emerged as the most robust gene sets for distinguishing senescence. HernandezSegura and SAUL_SEN_MAYO showed the strongest effect sizes and classification performance (**Supplementary Figure 5**), whereas SeneQuest and CellAge were less specific, especially in differentiating senescence from quiescence. In neurons, SeneQuest and CellAge displayed inverse trends (*i.e.*, senescence scores decrease from proliferative to senescent samples, compared to other cell types), possibly reflecting the unique, post-mitotic nature of neurons and their non-canonical, senescence-like phenotype (cell-cycle arrested but metabolically and secretorily active)(10,37). This suggests that SeneQuest and CellAge may be composed of proliferation-linked genes, whereas HernandezSegura and SAUL_SEN_MAYO capture broader senescence features. Notably, while we excluded gene sets exclusively representing the SASP due to its known context-dependence and variability, SAUL_SEN_MAYO includes a substantial number of genes predicted to be secreted (**Supplementary Figure 2A**). This suggests that it may capture a core SASP component relevant across cell types, potentially explaining its strong performance even in post-mitotic contexts. In contrast, HernandezSegura includes fewer secreted genes, suggesting that SASP alone does not fully account for the robustness of these gene sets in marking senescence.

Scores incorporate gene directionality, *i.e.*, whether genes are expected to be upregulated or downregulated in senescence (**Figure 2A**), which strongly influences their interpretation. For example, ignoring directionality in CellAge yielded higher senescence scores in proliferative than senescent samples and reduced classification accuracy (**Figure 3B**).

### Enrichment-Based Methods Enable Group-Level Interpretation

To evaluate the nine gene sets using an enrichment-based framework, we applied GSEA to gene rankings derived from linear models contrasting proliferative, quiescent, and senescent cellular states (see Methods).

Highlighting gene set members on volcano plots of differential expression (statistical significance vs. fold-changes, **Figure 4A**), summarising those models, enables visual assessment if genes within a phenotype-linked set consistently shift their expression according to expectation. CellAge, HernandezSegura, and SAUL_SEN_MAYO showed clear, consistent transcriptional differences between senescent and proliferative samples, while HernandezSegura and SAUL_SEN_MAYO also performed well in the senescent vs. quiescent contrast. SeneQuest, in line with score-based results, reflected non-proliferative features, such as a strong depletion of genes expected to be downregulated in senescent samples when compared with proliferative cells, but no clear signal in the senescence vs. quiescence contrast. Gene sets with annotated directionality and more variable differential expression statistics, such as CellAge and SeneQuest in the senescent vs. quiescent contrast, suggest a more context-dependent performance in marking the phenotype. Gene sets lacking directionality, such as GOBP_NEGATIVE_ and GOBP_POSITIVE_REGULATION_OF_CELLULAR_SENESCENCE, show weak enrichment and low fold changes, also suggesting context-dependent relevance rather than a global response of senescence.

**Figure 4.**
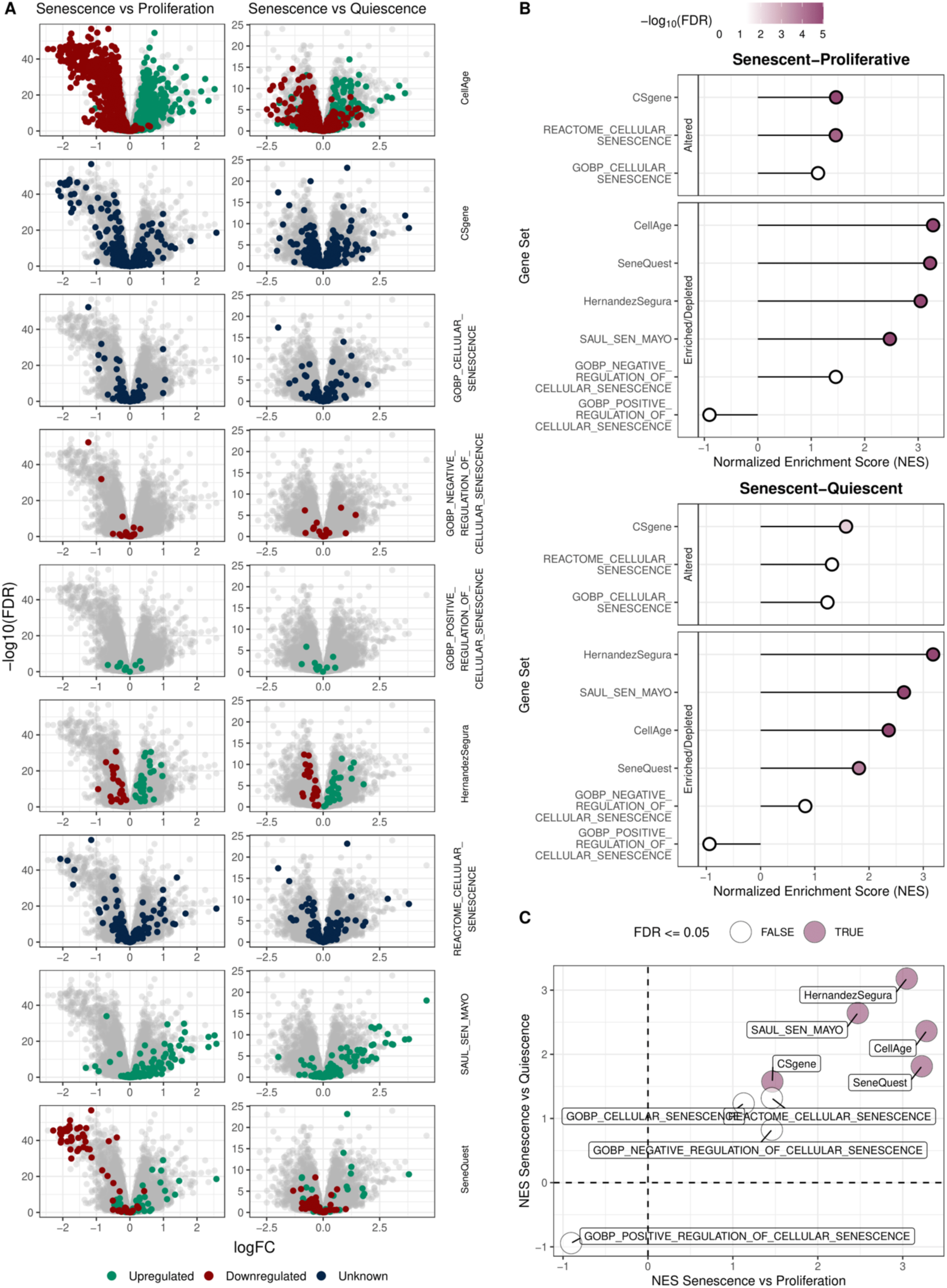
Performance of enrichment-based phenotype quantification method across gene signatures. (A) Volcano plots depicting differential gene expression results for the comparisons Senescent vs Proliferative (first column) and Senescent vs Quiescent (second column), across the nine gene signatures under analysis. Each dot represents a gene, with colour indicating its expected behaviour in senescence: red for downregulated, green for upregulated, blue for genes without directional annotation, and grey for genes not included in the given signature. (B) Gene Set Enrichment Analysis (GSEA) results for both comparisons involving senescence. Each plot summarises the Normalised Enrichment Scores (NES) for gene sets either lacking directional annotation (“Altered”) or annotated as “Enriched/Depleted” (see Methods, section 2.1.2). NES is shown on the x-axis, with gene sets on the y-axis, with a colour gradient proportional to the FDR. An FDR of above 0.05 is represented in white. (C) Scatter plot showing the NES from GSEA for each gene signature in two contrasts: Senescent vs. Proliferative (x-axis) and Senescent vs. Quiescent (y-axis). Purple dots represent signatures significantly enriched in both contrasts (FDR <= 0.05), while white dots indicate non-significance in at least one of the contrasts.

Gene Set Enrichment Analysis confirmed significant enrichment for CellAge, SeneQuest, CSgene, SAUL_SEN_MAYO and HernandezSegura across comparisons between the three cellular states (**Figure 4B, C**). Others, such as the GOBP and REACTOME gene sets, did not reach significance in the enrichment-based analysis for the two comparisons involving senescence.

### Exploring Best-Performing Senescence Gene Sets

Across score- and enrichment-based analyses, gene sets HernandezSegura, CellAge, SAUL_SEN_MAYO, and SeneQuest consistently emerged as robust indicators of senescence. These sets consistently discriminated senescent samples from both proliferative and quiescent cellular states across the considered metrics (**Figure 5A-B**). In contrast, MSigDB gene sets, despite their popularity in GSEA-based studies, performed poorly as senescence markers across most score- and enrichment-based metrics.

**Figure 5.**
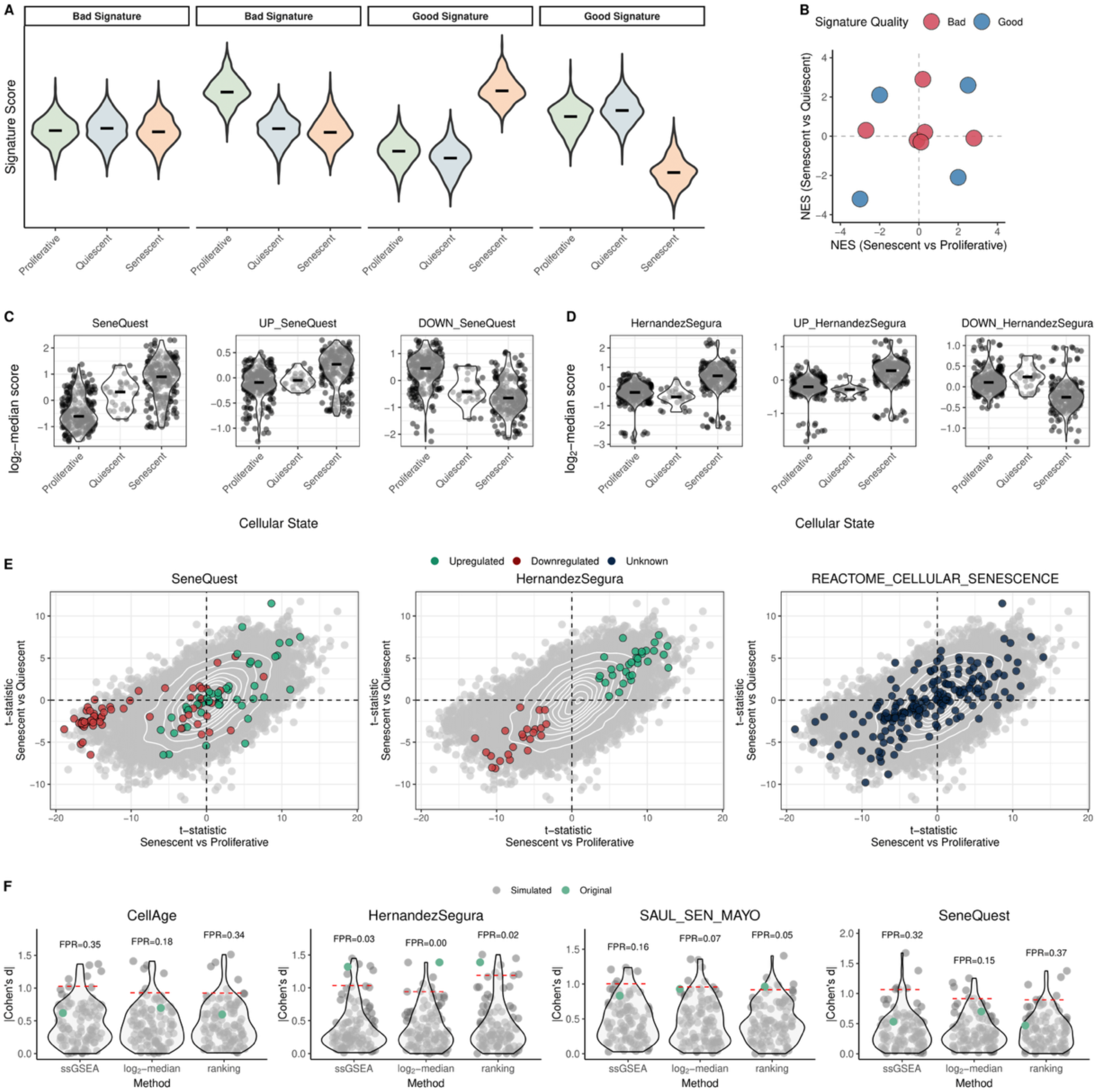
Comparative performance of senescence signatures. (A) Example data illustrating the behaviour of good and bad gene signatures using score-based methods. A good signature produces a score that separates senescent from not only proliferative, but also quiescent samples. Poor signatures show overlapping distributions between senescent and other conditions. (B) Scatter plot of GSEA NES for mock signatures. A good signature shows strong and consistent enrichment (positive or negative NES) in both contrasts (Senescent vs. Proliferative and Senescent vs. Quiescent), falling far from the NES = 0 axes. Poor signatures show weak or inconsistent enrichment across contrasts. Red and blue colours respectively indicate Bad and Good classifications based on NES strength. (C) Distribution of log_2_-median scores for CellAge and its UP- and DOWN-regulated subsets across proliferative, quiescent and senescent samples. The upregulated subset discriminates senescence from quiescence better, while the downregulated genes show less separation, indicating possible dominance by non-specific proliferation-associated genes. White-shaded density violins are overlaid to aid visualisation of distributions. (D) Distribution of log_2_-median scores for HernandezSegura and its up- and downregulated subsets across proliferative, quiescent and senescent samples, with both UP and DOWN compartments showing good discriminatory power between senescent and proliferative/quiescent samples. (E) Scatter plots of t-statistics of differential gene expression for the Senescent vs. Proliferative (x-axis) and Senescent vs. Quiescent (y-axis) comparisons. Each grey dot represents a gene, with signature genes highlighted for SeneQuest, HernandezSegura and REACTOME_CELLULAR_SENESCENCE (left to right). Colour indicates the expected behaviour of the genes in the set upon senescence: red for downregulated, green for upregulated, dark blue for genes without directional annotation, and grey for genes not included in the given signature. Density contour lines in white. (F) Violin plots showing the null distribution of absolute Cohen’s d effect sizes across the three scoring methods for 100 randomly simulated signatures. Each grey dot represents a simulated signature of the same size in number of genes as the original, while green represents the original tested signature. The dashed red line indicates the 95^th^ percentile of the null distribution. False positive rates (FPRs) are shown above each violin.

**SeneQuest**’s discriminatory power improved when its up- and down-regulated components were analysed separately. The putatively downregulated genes showed similar scores in senescent and quiescent samples (**Figure 5C**), indicating downregulation in senescence relative to proliferation but little distinction from quiescence, likely reflecting non-specific cell cycle arrest (**Figure 5E**). In contrast, the putatively upregulated genes effectively discriminated senescence from quiescence (**Figure 5C**), suggesting that they dominate the set’s performance in marking senescence. By comparison, both up- and down-regulated components of HernandezSegura effectively distinguish senescence from both proliferation and quiescence (**Figure 5D-E**). Conversely, as an additional example, REACTOME_CELLULAR_SENESCENCE shows poor value in discriminating between phenotypes, with its genes showing little differential expression between senescence and quiescence or proliferative samples (**Figure 5E, 3A**).

**CellAge** showed solid performance in marking senescence, particularly in enrichment based analyses, but in a cell-type- and stimulus-dependent manner. The scores observed in neurons may reflect a greater reliance on proliferation-associated programmes such as E2F targets, MYC signalling and mitotic checkpoint control (**Supplementary Figure 2B**), potentially limiting its applicability to non-dividing differentiated cells.

**HernandezSegura** emerged as the most consistent senescence-marking gene set, despite having few genes annotated as senescence-related in functional databases (**Figure 2A** and **Supplementary Figure 2B**). This suggests it captures genes with consistent expression changes upon senescence, although not necessarily the most extreme (**Figure 4A**), and therefore more likely to reflect a core transversal senescence signature, even if no functional link with the phenotype has been established yet.

**SAUL_SEN_MAYO**‘s robustness may stem from its enrichment in genes encoding secreted proteins (**Supplementary Figure 2A**), suggesting it captures core features of the senescence programme beyond cell cycle arrest, possibly driven by inflammation-related pathways such as JAK/STAT3 and NF-κB signalling (**Supplementary Figure 2B**).

When benchmarked against random gene sets for distinguishing senescence from quiescence, HernandezSegura and SAUL_SEN_MAYO showed low mean false positive rates (FPR, 0.02 and 0.09). In contrast, SeneQuest and CellAge showed higher mean FPRs (0.28 and 0.29), suggesting that they are the top markers of cell cycle arrest but not specifically of senescence (**Figure 5F, Supplementary Figure 6**).

### Trade-Offs

To assess the robustness of score- and enrichment-based approaches, we evaluated their performance as markers across varying sample sizes (**Figure 6A**) and gene-set sizes (**Figure 6B**). As expected, the size of gene sets is relevant for their performance, as each gene potentially brings to the signature additional information for the discrimination between phenotypes and redundancy to the collective marking ability. The CellAge signature (1259 genes) more strongly discriminates proliferative from senescent samples than Hernandez-Segura (55 genes) (**Figure 6A**), and random gene sampling also shows proportionality between gene set size and marking performance, whatever the signature and the marking approach (**Figure 6B**). However, this association appears to only hold up to a certain set size, likely the minimum number of genes comprising the full non-redundant marking information. This number depends on the signature and the marking approach. If GSEA is used, only about one third of the CellAge genes are needed for its optimal performance (better than the full signature’s) in discriminating proliferative from senescent samples (**Figure 6B**, top left). With the log_2_-median score, just over 100 genes are needed for a performance similar to the full signature’s (**Figure 6B**, right). The full Hernandez-Segura set is needed for maximum performance with GSEA (**Figure 6B**, left) but less than 20 genes are required for peak performance with a score-based approach, with this performance being superior to CellAge’s for the same number of genes (**Figure 6B**, right insets). This suggests that the sparser Hernandez-Segura signature contains a few highly informative genes for the proliferative vs. senescent contrast not included in CellAge.

**Figure 6.**
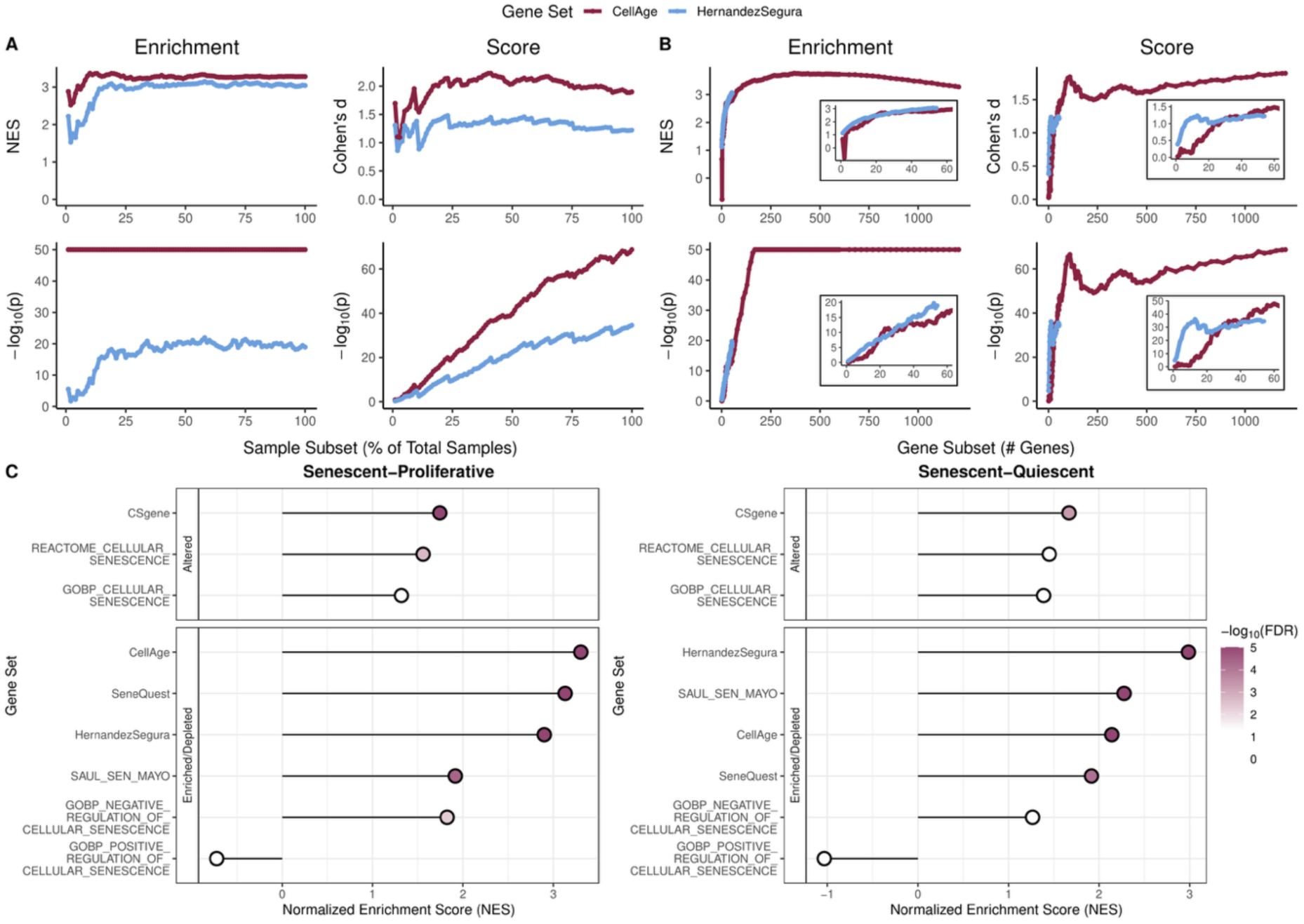
Performance of score- and enrichment-based approaches under varying sample and gene set sizes. (A) Effect sizes (top row) and p-values (bottom row), for GSEA (left “Enrichment” column) and log2-median score (right “Score” column), for pairwise comparisons between proliferative and senescent samples over random subsets of samples of increasing size (2-100% of 240 proliferative and 272 senescent samples, resampling for each percentage). (B) Effect sizes and p-values for the same methods in (A) over random gene subsets of increasing size from each representative gene set (one-gene increments up to 55, the Hernandez–Segura size, followed by 10-gene (starting at 69 genes) and then 30-gene (starting at 599 genes) steps up to the CellAge size, resampling genes for each number). Both panels show representative gene signatures: “small” (55 genes) HernandezSegura (blue) and “large” (1259 genes) CellAge (bordeaux). “Zoom-in” insets show the same results for up to 60-gene subsets. (C) Lollipop plots of GSEA results for all nine gene signatures using a subset of the data consisting only of proliferative, quiescent and radiation-induced senescent samples - mirroring the format of Figure 4B. In lollipop plots, NES are shown for both altered and enriched/depleted gene sets, with NES on the x-axis and gene sets on the y-axis, coloured by the FDR.

Our analyses show that less than 50 samples of each phenotype are enough for the signatures’ maximum performance (**Figure 6A**). With GSEA, that plateauing happens with even fewer samples for gene-rich CellAge (**Figure 6A**, top left). As expected due to the approaches’ nature, significance of GSEA-based results is proportional to the number of genes and saturates when that number gets into the hundreds (**Figure 6A,B**, bottom left), whereas as significance of score-based results follows the proportionality with sample size that is typical of testing differences between pairs of distributions when the effect size does not change (**Figure 6A**, bottom right).

The performance as markers of certain gene sets, profiled by enrichment methods, was influenced by the choice of senescence inducer. When limiting our analyses to proliferative, quiescent and radiation-induced senescence samples, they (*e.g.*, GOBP_NEGATIVE_REGULATION_OF_CELLULAR_SENESCENCE) marked senescence more strongly than when all senescent subtypes were included (**Figure 6C**). This suggests that some gene sets are inducer-specific signatures and underperform in other senescence contexts. The per-sample resolution of graphical depictions of distributions of scores allows to visually capture such context-specific trends (**Supplementary Figure 4**).

### From Biological Insight to Translational Potential

Senescence has been linked to multiple diseases (38), so quantitatively marking it in large cohorts of samples could reveal associations with clinical features. Given the robustness of the HernandezSegura and SAUL_SEN_MAYO gene sets, we used the discovery mode of `markeR` to test their potential to mark senescence in human tissues GTEx data across 49 healthy tissues (39). While there is no definitive ground truth for relative senescence levels in these tissues, it is generally accepted that senescent cells accumulate with age (36). Thus, if these gene sets are effective, we expect their scores or enrichments to positively correlate with age. HernandezSegura is shown here as the top-performing gene set from the benchmarking analysis (including the lowest mean FPR in distinguishing senescence from quiescence; **Figure 5F**, **Supplementary Figure 6**); homologous results for SAUL_SEN_MAYO are shown in **Supplementary Figures 7** and **8**.

Each tissue exhibited a distinct median senescence score (**Figure 7A**). Consistently, from the *in vitro* senescence datasets, we observed that baseline senescence scores can differ between cell types; for instance, proliferative mesenchymal cells can score higher than senescent melanocytes (**Figure 7B**). This underscores how tissue heterogeneity can dilute senescence signatures. Therefore, score differences between tissues (**Figure 7A**) may reflect distinct cell-type compositions rather than differences in senescence levels *per se*.

**Figure 7.**
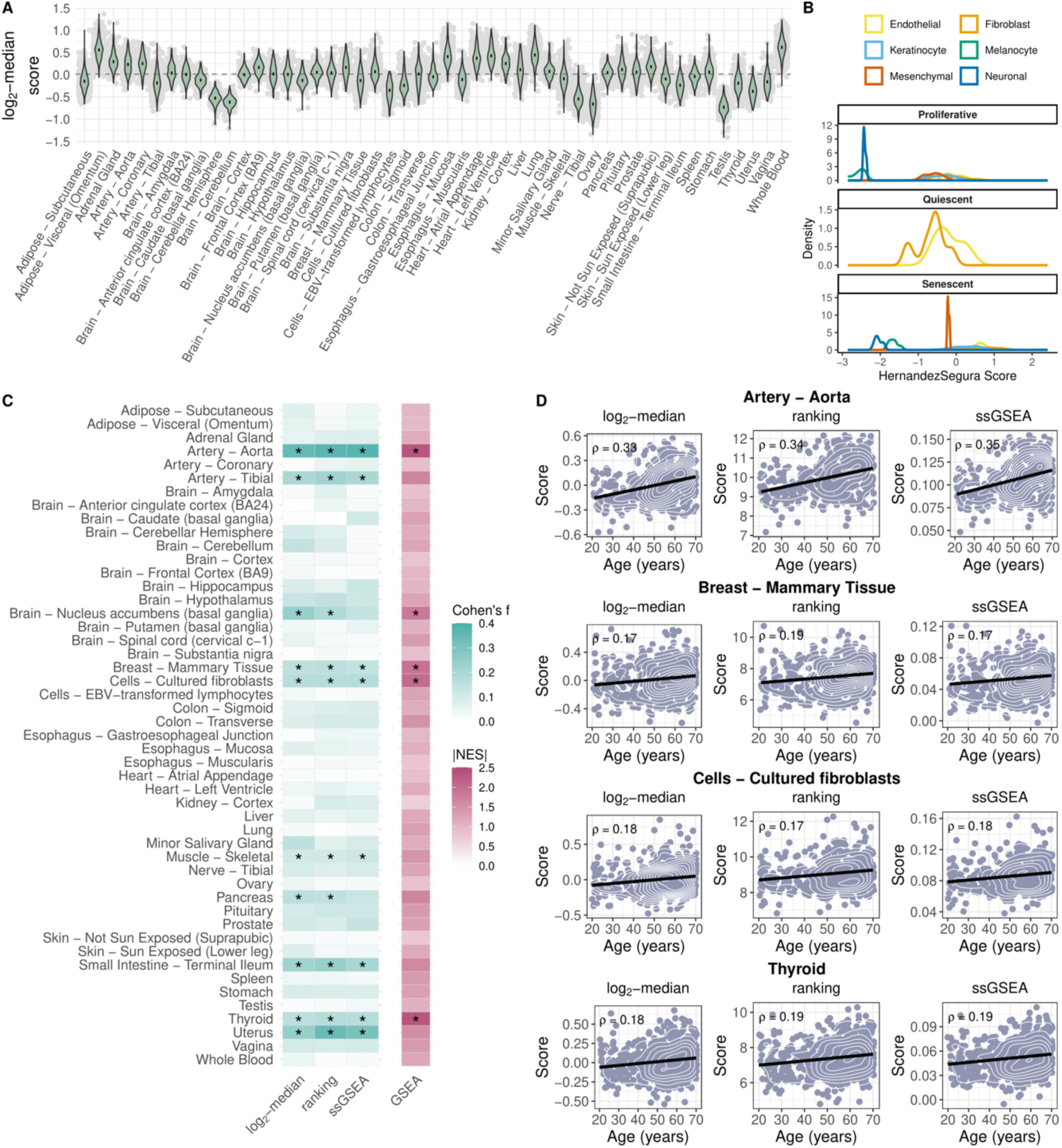
Influence of age on HernandezSegura senescence signature scoring and enrichment across human tissues. (A) Violin plots of distribution of senescence scores across 49 tissues from GTEx v8, using the log2-median method and the HernandezSegura signature as an example. Each grey dot represents a sample, and the tissues are ordered alphabetically along the x-axis. (B) Distributions of senescence scores in proliferative, senescent and quiescent samples from in vitro datasets show that, for certain cell types, some proliferative samples display higher scores than senescent samples of other cell types. This highlights the strong cell-type specificity and variable baseline (i.e., in proliferative samples) levels of senescence signature scores. (C) Heatmap summarising the association between donors’ age and senescence scores across GTEx tissues. Rows represent tissues and columns represent scoring-based approaches (log2-median, ranking and ssGSEA) using Cohen’s f as a metric of variable association (white to green), and GSEA-based enrichment analysis (NES, white to bordeaux) using age as the variable. Asterisks denote significant associations (FDR ≤ 0.05). For the score-based approaches, p-values were obtained from 1,000 random permutations of the age variable, whereas for GSEA, the p-value is derived from the method’s internal enrichment testing procedure. (D) Scatter plots showing score vs age for the four tissues with significant associations across all four methods in (C). Each row corresponds to one tissue, with plots for the log2-median, ranking and ssGSEA methods. The age in years (x-axis) is jittered for donor privacy. Black linear regression line and Spearman’s correlation coefficient π are shown. Density contour lines in white aid visualisation of dot distributions. Only genes detectable across all 49 tissues were retained for this analysis; the impact of this filtering strategy is shown in **Supplementary** Figure 11.

For HernandezSegura, only four tissues (Aorta, Mammary Tissue, Cultured Fibroblasts, and Thyroid) showed significant age-related changes in both score and enrichment approaches (**Figure 7C, D**). Cultured Fibroblasts are of particular interest, given that fibroblasts were key in the derivation and validation of the signature. The observation of an age-related increase in the signature’s strength therein, despite the large inter-individual variability, is compatible with the hypothesis of an accumulation of senescent cells with age (**Figure 7D**). The strongest signal observed for the aorta is also consistent with links between senescence and age-related vascular dysfunction (40). Unlike more complex organs, the aorta has low cellular diversity and is mainly composed of endothelial and vascular smooth muscle cells (41). Its lifelong exposure to mechanical strain (42) may also contribute to the observed age-related changes. However, these interpretations should be made with caution. On one hand, a similar pattern was not seen in other arterial tissues, such as the tibial. On the other, we did not detect significant age-related strengthening of the senescence signature in tissues such as the skin, lung or pancreas, despite their known links to age-related senescence (11). We could not detect it either in two independent datasets of tissues in which age-related senescence accumulation would be expected: inner forearm epidermis and whole lung homogenate (11) (**Supplementary Figure 9**).

SAUL_SEN_MAYO includes many SASP genes, including inflammatory ones, so results may partly reflect age-related inflammation (inflamm-ageing) rather than senescence (44). Using the same criteria as for HernandezSegura, eight tissues showed age-related alterations in scores or enrichment (**Supplementary Figure 7**), including brain regions such as the Anterior Cingulate Cortex and Hippocampus. In some tissues (*e.g.*, Sigmoid Colon, Tibial Nerve), the score decreased with age, contrary to what was expected (**Supplementary Figure 7B**). To assess the potential role of inflammation in our observations, we split the gene set into inflammation-related (N = 81) and -unrelated (N = 44) genes. Both subsets performed similarly in the *in vitro* senescence datasets, confirming their robustness in that context with a ground truth for senescence (**Supplementary Figure 8A-B**). In GTEx, the association of the inflammatory and non-inflammatory gene subsets with age varies across tissues (**Supplementary Figure 8C**). The absence of a consistent trend across tissues indicates that the observations made for SAUL_SEN_MAYO cannot be solely attributed to inflammageing, and suggest a variable combination of the signals of senescence and inflammation.

Overall, interpretation of changes in molecular signatures in tissue data must take into account that cellular composition changes with age (*e.g.,* more adipose and less epithelial cells in the breast, more granulocytes in blood)(45), and GTEx samples are cross-sectional, not longitudinal (which, while ideal, are impractical). To control for cell type composition, we reran the enrichment analysis incorporating estimated cell type scores as covariates (see Methods; **Supplementary Figure 10**, HernandezSegura used as an example). NES values and their percentile ranks relative to a null distribution were highly consistent between the two models across all 49 tissues, with only minor fluctuations. This concordance likely reflects the inherently weak transcriptional signal of senescent cells, as senescent cells are expected to represent only a small minority of cells in functional tissues, such that compositional adjustment has limited leverage on an already dilute signal. The large inter-individual variability, inherent to a cross-sectional cohort such as GTEx, may further mask meaningful senescence-specific signals in bulk transcriptomic data.

### markeR as a holistic framework for the evaluation of gene sets as phenotypic markers

A central finding from this work is that the choices of gene set and quantification method shape the biological interpretation of the outcomes of markeR’s usage. The senescence case study demonstrates this directly: HernandezSegura and SAUL_SEN_MAYO emerged as robust markers across diverse cell types and stressors, while commonly used MSigDB sets performed poorly and others such as CellAge showed marked context-dependence.

This challenge extends beyond senescence, as any phenotype lacking a universal marker faces the same problem: multiple competing gene sets and quantification strategies are available, with no principled framework guiding their evaluation. While existing tools address individual aspects of this problem, including scoring methods (*e.g.*, singscore (46), UCell (47), GSVA (48), irGSEA (49)), enrichment-based tools (*e.g.*, GSEA/fgsea (50,51), decoupleR (52)), benchmarking frameworks (*e.g.*, GSEABenchmarkeR (53)), and phenotype-specific classifiers (*e.g.*, hUSI (54), SenePy (55)), rigorously evaluating a gene set as a phenotypic marker currently requires independently combining several of these, each with its own inputs, assumptions, and learning curves. markeR was designed to address this directly: by unifying multiple quantification strategies within a single modular framework, it allows researchers to immediately assess whether the marking ability of gene sets is consistent across methods, while providing supporting evidence such as individual gene contributions, empirical null benchmarking, and effect size metrics, all from standard transcriptomic inputs (*i.e.*, a gene expression matrix, a gene set list, and sample metadata) (**Table 1**).

**Table 1.**
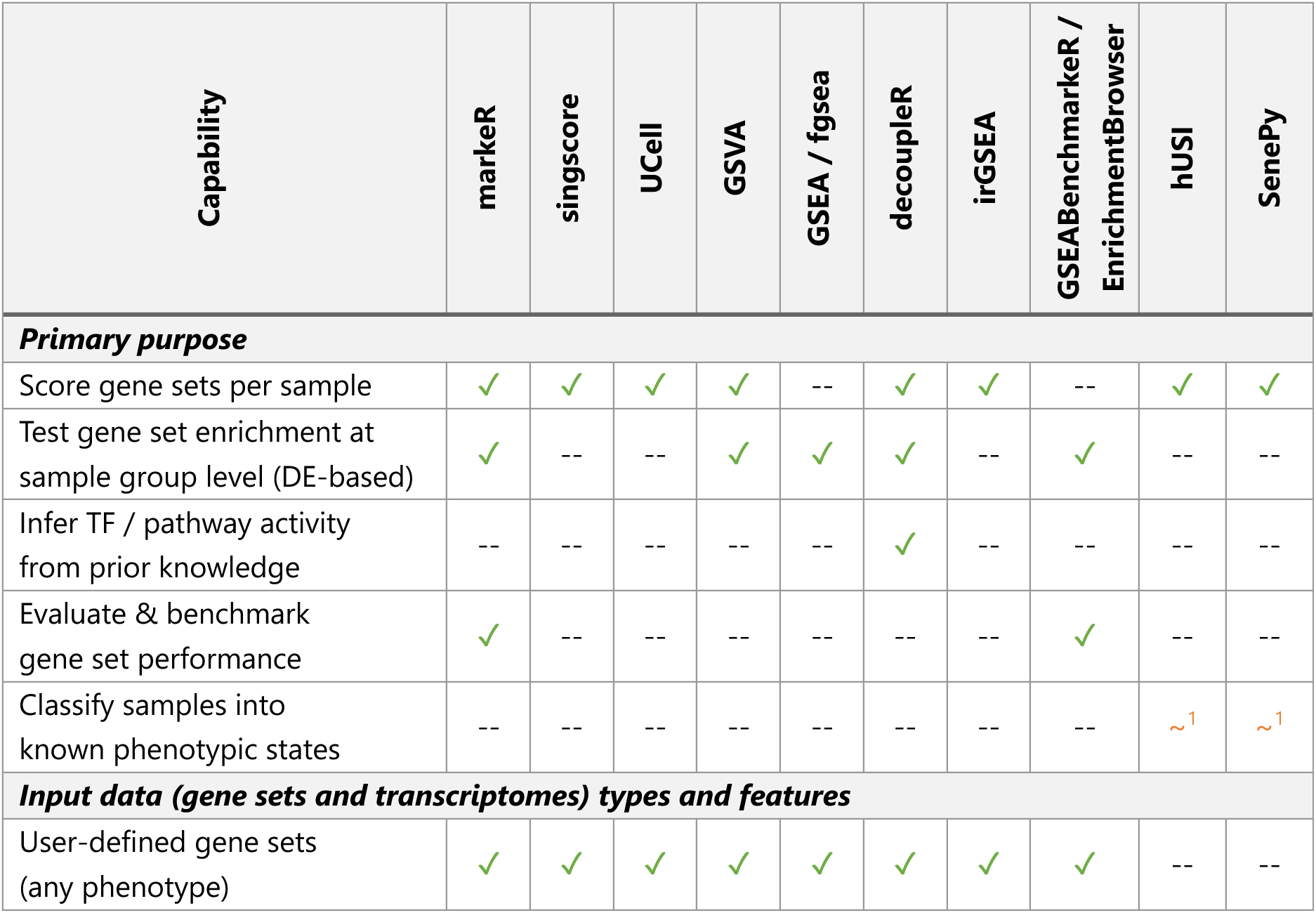
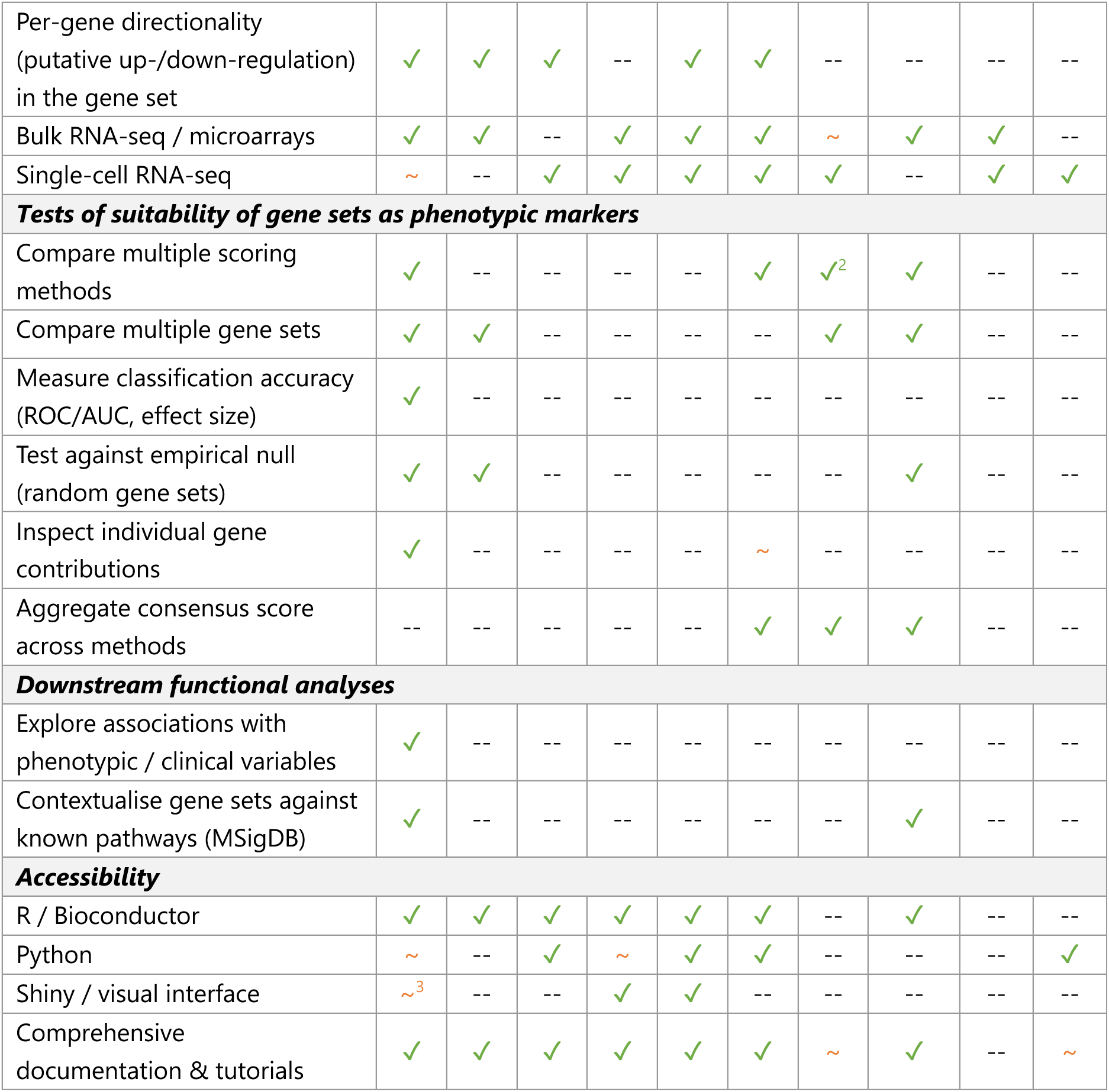
Gene set analysis tools (with scoring and enrichment strategies) differ substantially in their purpose, scope, and analytical capabilities. This table compares markeR with existing tools across five user-centred dimensions: primary analytical purpose, supported input data features, capabilities for testing gene sets as phenotypic markers, functional analyses, and accessibility. The first eight tools are applicable to any user-defined phenotype; hUSI and SenePy offer curated, pre-trained signatures for senescence. ✓ full support; ∼ partial or planned support; -- not supported. ^1^ hUSI and SenePy compute senescence-associated scores that are enriched in senescent relative to non-senescent samples, rather than providing binary sample classification. Neither is designed as a general-purpose framework for user-defined phenotypes. Support is therefore marked as partial (∼). ^2^ While irGSEA also integrates multiple scoring methods (AUCell, UCell, singscore, ssGSEA, JASMINE, Viper), it aggregates results to yield a consensus score. In markeR, however, disagreement across methods is explicitly shown to help users assess the robustness of their findings. ^3^ Shiny / visual interface: markeR Shiny app is under development and planned for a future release.

markeR’s modular design furthermore enables integration of new quantification methods as they emerge, and its applicability extends to any phenotype with a hypothesised gene expression signature: from senescence to sex-specific gene expression, acute respiratory viral response, cellular proliferation, and environmental stress responses in *Saccharomyces cerevisiae* (**Supplementary Figure 12**).

## DISCUSSION

Quantifying complex cellular phenotypes like senescence in transcriptomic samples often requires integrating multiple gene signatures and analytical strategies, as no single gene functions as a universal marker. To meet such need, we developed `markeR`, a software for flexibly analysing and evaluating gene sets as transcriptomic phenotypic markers.

Key strengths of `markeR` are its modular architecture, supporting the integration of new functionalities (e.g., enrichment methods beyond GSEA), and its emphasis on continuous, quantitative metrics over binary classification methods, reflecting the non-discrete nature of biological phenotypes. Rather than enforcing strict cut-offs, `markeR` encourages interpreting scores in the context of their distribution, allowing more nuanced insights, particularly for cases deemed borderline, and supporting flexible, data-informed decisions. A Shiny-based interface is planned to improve accessibility and integration with public dataset repositories (*e.g.*, GEO (27)), and upcoming releases will implement standardised metrics for cross-cell-type comparisons (23). While senescence serves as the primary case study herein, markeR is applicable to any phenotype characterizable by gene sets (**Supplementary Figure 12**).

Applying `markeR` to marking senescence yielded both technical and biological insights. Scoring methods offer sample-level resolution but are sensitive to the behaviour of individual genes, potentially skewing interpretability when, for instance, the goal is to evaluate an entire pathway as a phenotypic marker. Enrichment-based methods focus on coordinated gene behaviour but are limited to sample group-level comparisons. Even among scoring methods, applicability tradeoffs exist: rank-based approaches (*e.g.*, ssGSEA and rank scoring) tend to be more robust to noise, while magnitude-based methods (*e.g.*, log_2_-median) may better detect expression shifts in technically well-controlled settings. Ultimately, these methods should be seen as complementary and applied in an integrative manner, guided by the same biological question.

No gene set performed optimally in specifically marking senescence across all cell types analysed herein. However, HernandezSegura and SAUL_SEN_MAYO stood out for their consistency across metrics and low false positive rates, suggesting they capture core senescence molecular features. In contrast, commonly used senescence-associated MSigDB gene sets performed poorly, and others, like CSGene and SeneQuest, have their performance affected by oblivion to the direction of marker gene expression changes upon phenotype and enrichment in non-specific proliferation-related genes, respectively.

Context-dependence emerged as a major, though expected challenge. Senescence is not a discrete state but a complex phenotype influenced by inducing stimulus, cell type and microenvironment (10). In human tissues, this complexity is reflected in the inter-individual and often age-related variability in cell type composition (45), and its study is hampered by the typically low abundance of naturally occurring senescent cells. Although we used quiescent samples as non-proliferative controls, other non-senescent states (*e.g.*, differentiated cells), may still affect marker specificity.

Some tissues, like cultured fibroblasts and the aorta, showed association between molecular signals of senescence and donor’s age, and could be seen as positive controls for the accumulation of senescent cells with ageing. However, defining ’positive controls’ for senescence is not trivial. Tissues like skin, lung or pancreas are often cited for accumulating senescent cells with age, based on markers like p16^(INK4a)^ or p21^(CIP1/WAF1)^ (11,56). Yet these markers are not exclusive to senescent cells. For example, macrophages can express p16^(INK4a)^ in non-senescent inflammatory contexts (57), and p21^(CIP1/WAF1)^ is a stress-responsive gene that does not necessarily indicate irreversible growth arrest. Thus, some reported age-associated ’senescence’ signals may reflect immune or stress responses, rather than senescence itself. In this context, the lack of significant age associations in some tissues, despite previous reports, likely reflect the limited specificity of senescence markers and the inability of bulk RNA-seq to resolve cell-type-specific processes. Moreover, GTEx data are cross-sectional rather than longitudinal and marked by the expectedly large natural inter-individual variability, further complicating interpretation. These findings emphasise the need to interpret senescence molecular signatures with caution when translating them from controlled *in vitro* settings to the complex cellular environments of human tissues. Cell type composition is the only confounder formally addressable in this data, and we show that the observed age-senescence associations do not change after this correction. The rarity of senescent cells in functional tissues and the inter-individual differences in senescence burden inevitably dilute the senescence transcriptional signal in bulk transcriptomes, underscoring that the age-senescence associations reported here should be regarded as exploratory, and studies combining orthogonal senescence markers with single-cell transcriptomics are needed to validate them.

That limitation illustrates the difficulty of establishing a ground truth for benchmarking and effective validation of robustness and universality of senescence gene signatures. The gene expression profiles used in this study are from cultures with evidence for senescence induction based on SA-β-galactosidase activity and p16^(INK4a)^ or p21^(CIP1/WAF1)^ (among others) mRNA or protein assays. However, none of these is a universal senescence marker and the respective assays involve cell populations and cannot resolve how many or which individual cells in each sample are senescent, with the sample’s gene expression profile inevitably reflecting contributions from non-senescent cells. Purifying senescent cell populations with flow cytometry is hampered by the absence of a universal surface marker. These constraints are not specific of markeR but reflect a fundamental unmet need in the field that markeR aims to help addressing.

Senescence is best understood as a heterogeneous collection of phenotypes shaped by the initiating stimulus, cell type and microenvironment, rather than as a single, uniform cellular state. This mirrors the current view of cancer as a spectrum of distinct yet overlapping transcriptional states, broadly defined by uncontrolled proliferation, rather than as a single, fixed phenotype. In contrast, the study of immune cell biology often emphasises unprecedent resolution at the level of specific, narrowly defined cell types. Applying such granular and rigid compartmentalisation to a systemic process like senescence risks overlooking broader molecular patterns and conserved mechanisms that transcend cell-type- and stimulus-specificity. A hybrid approach that recognises and integrates both the conserved hallmarks and the context-specific features of senescent cells is essential.

Quantifying senescence in biological samples from their bulk transcriptomes remains challenging, as molecular signals from rare senescent cells are diluted therein. While markeR is currently designed for bulk transcriptomes, several of its modules are, in principle, directly applicable to scRNA-seq data without modification. The log_2_-median-centred score, for example, can be applied per cell as-is, and is already implemented in scRNA-seq workflows such as scStudio (58). The individual gene module, including ROC/AUC analysis and PCA, is equally well-suited for single-cell data. For rank-based and ssGSEA scores, technical dropouts introduce many undetected genes that generate ties in rankings, potentially biasing scores and making them uninformative (59). A principled mitigation, as implemented in tools such as UCell (47), is to normalise against a reduced gene universe rather than the full transcriptome, down-weighting the contribution of undetected genes. For enrichment-based analyses, limma has demonstrated robust performance in scRNA-seq differential expression benchmarking (60), supporting its applicability in this context. Extending markeR with algorithms natively tailored for scRNA-seq remains part of our future plans.

Meanwhile, controlled *in vitro* datasets remain indispensable for benchmarking gene sets and workflows, and for mapping the transcriptional diversity of senescence across cell types, stimuli, and other specific microenvironmental contexts. These models will therefore continue to play a central role in refining our understanding of the molecular landscape of senescence.

## ACKNOWLEDGEMENTS

We thank the present and past members from the Disease Transcriptomics Lab, for the fruitful discussions, for testing the package’s functionalities during its development, and for providing valuable feedback on the manuscript. The authors used OpenAI’s GPT-4 model (accessed via ChatGPT) to assist with language polishing. The model was not used for content generation or data analysis. As proud ’research parasites’ (61), we processed publicly available datasets; we thank the researchers whose studies provided the data used in this work, for generating and sharing it, which proved invaluable to the feasibility of our study.

## AUTHOR CONTRIBUTIONS

Rita Martins-Silva: Conceptualisation, Formal analysis, Methodology, Writing – original draft, Data curation, Software, Visualisation. Alexandre Kaizeler: Methodology, Writing – original draft. Nuno L. Barbosa-Morais: Conceptualisation, Supervision, Writing – original draft.

## SUPPLEMENTARY DATA

##Supplementary Data are available at NAR online.

## CONFLICT OF INTEREST

None declared.

## FUNDING

This work is funded by national funds through Fundação para a Ciência e a Tecnologia, I.P. [UIDP/50005/2020 ; UI/BD/153368/2022 to R.M.-S. ; UI/BD/153363/2022 to A.K. ; CEECIND/00436/2018 to N.L.B.-M.].

## DATA AVAILABILITY

No data has been generated for this manuscript. The senescence compendium was compiled using publicly available RNA-seq datasets accessed under the following accession codes: E-MTAB-5403 (13), E-MTAB-9714 (62), GSE175533 (63), GSE60340 (64), GSE63577 (65), GSE64553 (66), GSE214410 (67), GSE222400 (68), GSE224070 (69), GSE230181 (70), GSE235768 (71), GSE247831 (72), GSE74324 (73), GSE75643 (74), GSE94928 (75), GSE130727 (76), GSE250224 (77), GSE196724 (78), GSE206677 (79), GSE217718 (80), GSE110268 (81), GSE213323 (82), E-MTAB-10969 (83). Data from The Genotype-Tissue Expression (GTEx) v8, already aligned, processed (filtered and normalised) and corrected for batch effects, was obtained from the GitHub repository associated with the Schneider et al., 2024 publication (https://github.com/DiseaseTranscriptomicsLab/voyAGEr) (33). Specific donor metadata, including the exact age of each donor, was retrieved through the dbGaP accession number phs000424.v9.p2.

The complete source code for the markeR package (v1.0.0), used throughout this manuscript, is publicly available through Bioconductor (doi : 10.18129/B9.bioc.markeR) and GitHub (https://github.com/DiseaseTranscriptomicsLab/markeR). The package can be installed in R using the command BiocManager::install(“markeR”). Python users can access the full markeR functionality via utility scripts based on the rpy2 bridge, available in the python/ directory of the GitHub repository. Additionally, the code for data preprocessing used in the senescence case study, along with the scripts to generate the figures in this paper, as well as the processed senescence datasets and gene sets used for the main figures, can be found in the paper branch of the markeR GitHub repository, as well as in FigShare (https://doi.org/10.6084/m9.figshare.31028227).

## Footnotes

1 Andrews, S. (2010). FastQC: A Quality Control Tool for High Throughput Sequence Data [Online]. Available online at: http://www.bioinformatics.babraham.ac.uk/projects/fastqc/

## SUPPLEMENTARY FIGURES

**Supplementary Figure 1.**
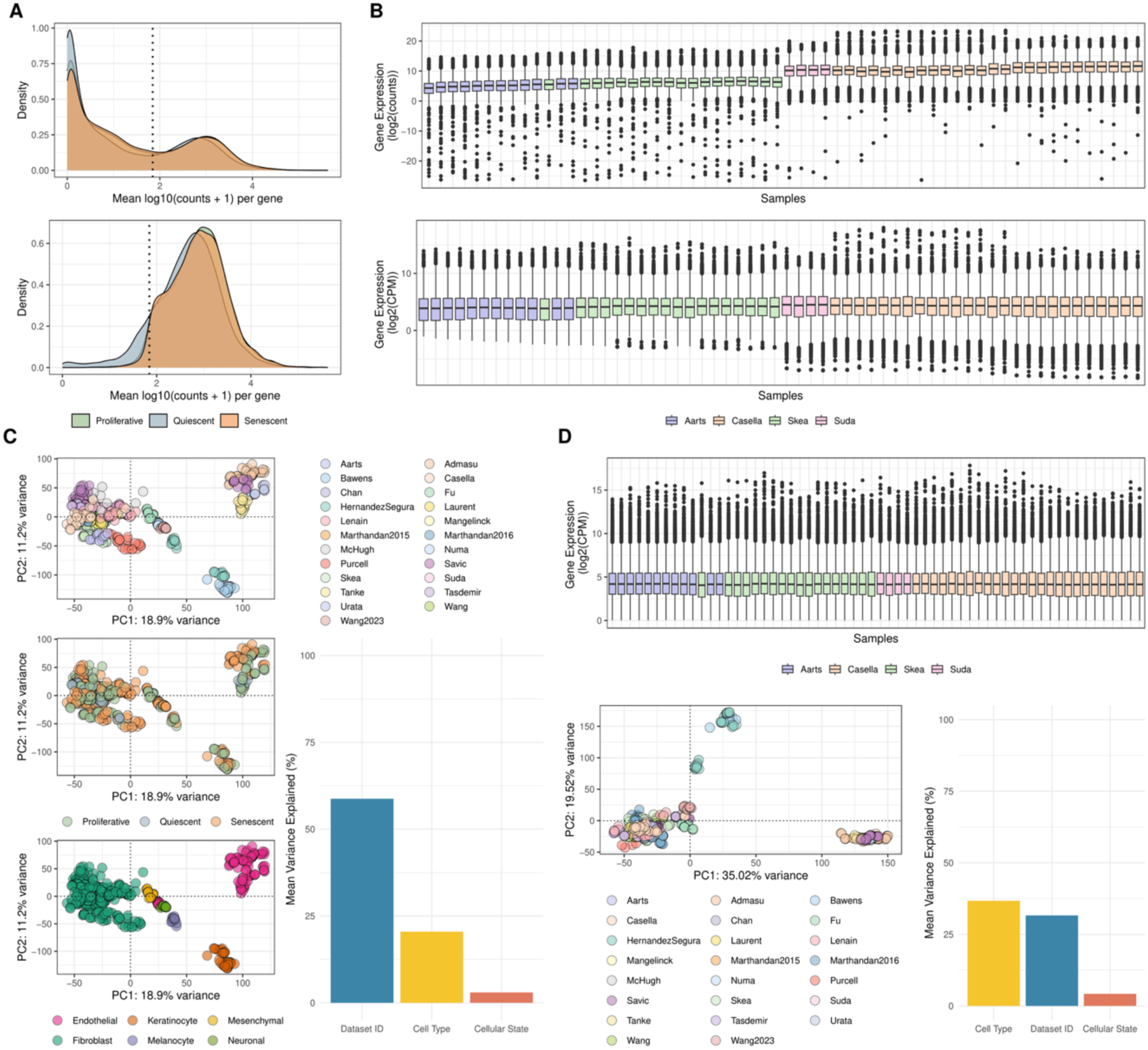
Pre-processing of senescence datasets. A total of 25 RNA-seq datasets comprising 545 samples from six human cell types were combined into one dataset and processed jointly. (A) Density plots of distributions of mean log10-transformed raw gene expression, with samples grouped by cellular state, before (top) and after (bottom) filtering. Lowly expressed genes were filtered out by retaining only those with an average of >70 reads in at least one condition (dashed line). (B) Box plots of log2CPM gene expression distributions across the 30 samples with the lowest and highest total read coverage. The top plot shows raw, filtered data, while the bottom shows data after normalisation, resulting in comparable median read counts across samples. (C) Principal Component Analysis (PCA) plot of filtered and normalised gene expression, with samples coloured by dataset ID (top left), cell type (top right), and cellular state (bottom left). (Bottom right) Bar plot of percentages of gene expression variance explained by those variables. (D) (Top) Box plots of log2CPM gene expression distributions across the same 30 samples as in (B) after batch correction, with no major inter-sample differences detected. (Bottom) The PCA plot of gene expression (left) and the bar plot of percentages of gene expression variance explained by the main variables after batch correction suggest its success, with dataset ID no longer dominating variance.

**Supplementary Figure 2.**
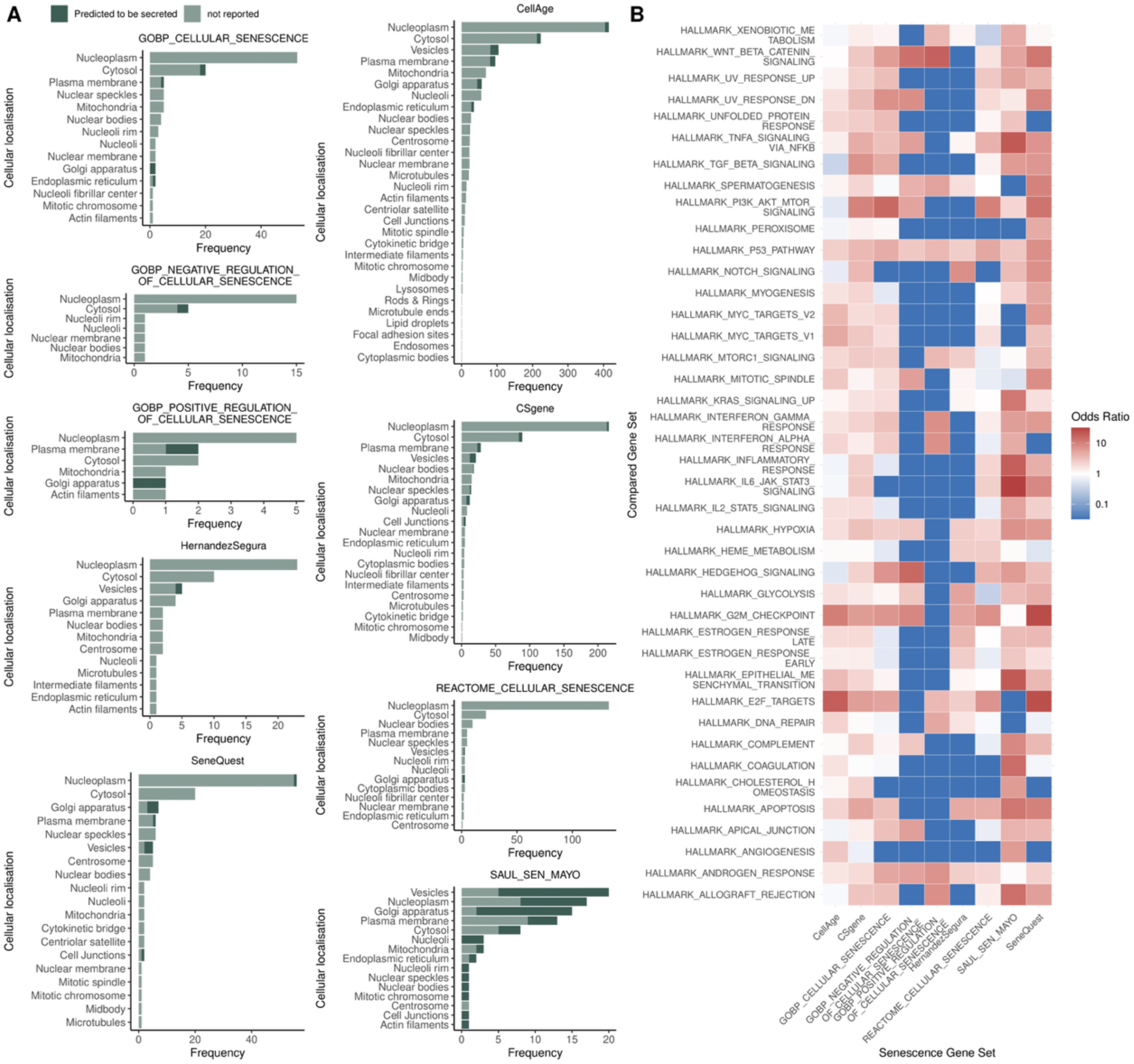
Subcellular localisation and functional enrichment of senescence gene signatures. (A) Cellular localisation of proteins encoded by genes in each set, based on annotations from the Human Protein Atlas. Bars represent the frequency of genes associated with each subcellular compartment. Genes predicted to be secreted are shown in dark green, while those with no reported extracellular localisation are shown in grey. Genes annotated to multiple compartments counted once per location. (B) Functional enrichment analysis (signature similarity feature of markeR) of the same gene sets using Hallmarks from MSigDB, chosen as the least redundant and most interpretable collection of biological processes therein. Heatmap of odds ratios (OR) of enrichment of genes from senescence signatures (columns) in Hallmarks (rows). The gene background consisted of all genes annotated in The Human Protein Atlas.

**Supplementary Figure 3.**
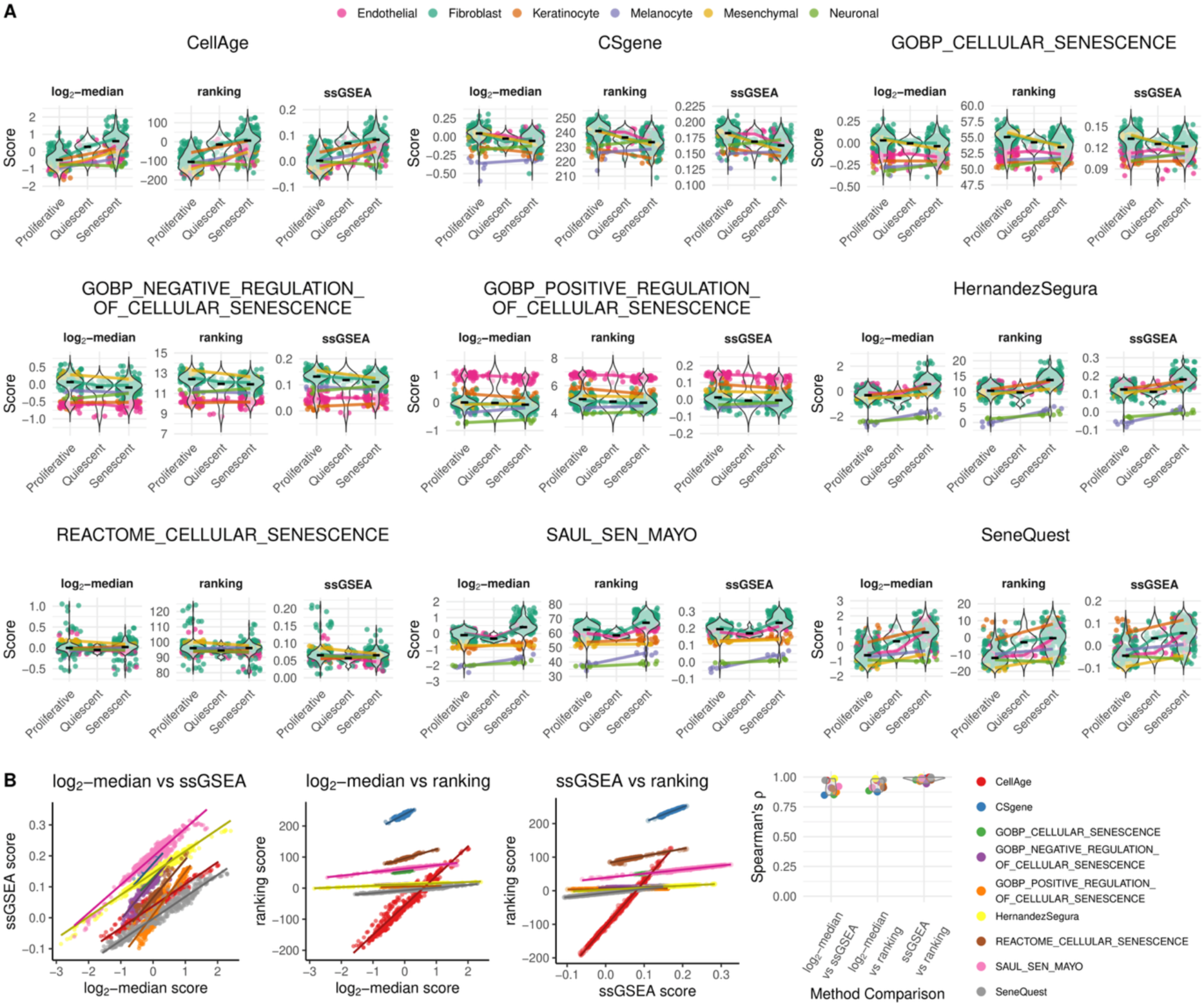
Performance and pairwise correlation of score-based phenotype quantification methods across gene signatures. (A) Violin plots showing the distribution of senescence scores for each gene signature, computed using three scoring methods: log_2_-median (left), ranking (middle), and ssGSEA (right). Each plot depicts the score distribution across sample types (proliferative, quiescent, senescent), coloured by cell type. Median scores per condition for each cell type are connected by coloured lines. (B) Pairwise comparisons of senescence scores across methods. The first three panels show scatter plots of log_2_-median vs. ssGSEA, log_2_-median vs. ranking, and ssGSEA vs. ranking. Dots (i.e., samples) are coloured by the gene set scored therein, with an associated regression line. The last panel shows the distribution of Spearman’s correlation coefficient ρ across all method pairs, confirming consistently strong correlations.

**Supplementary Figure 4.**
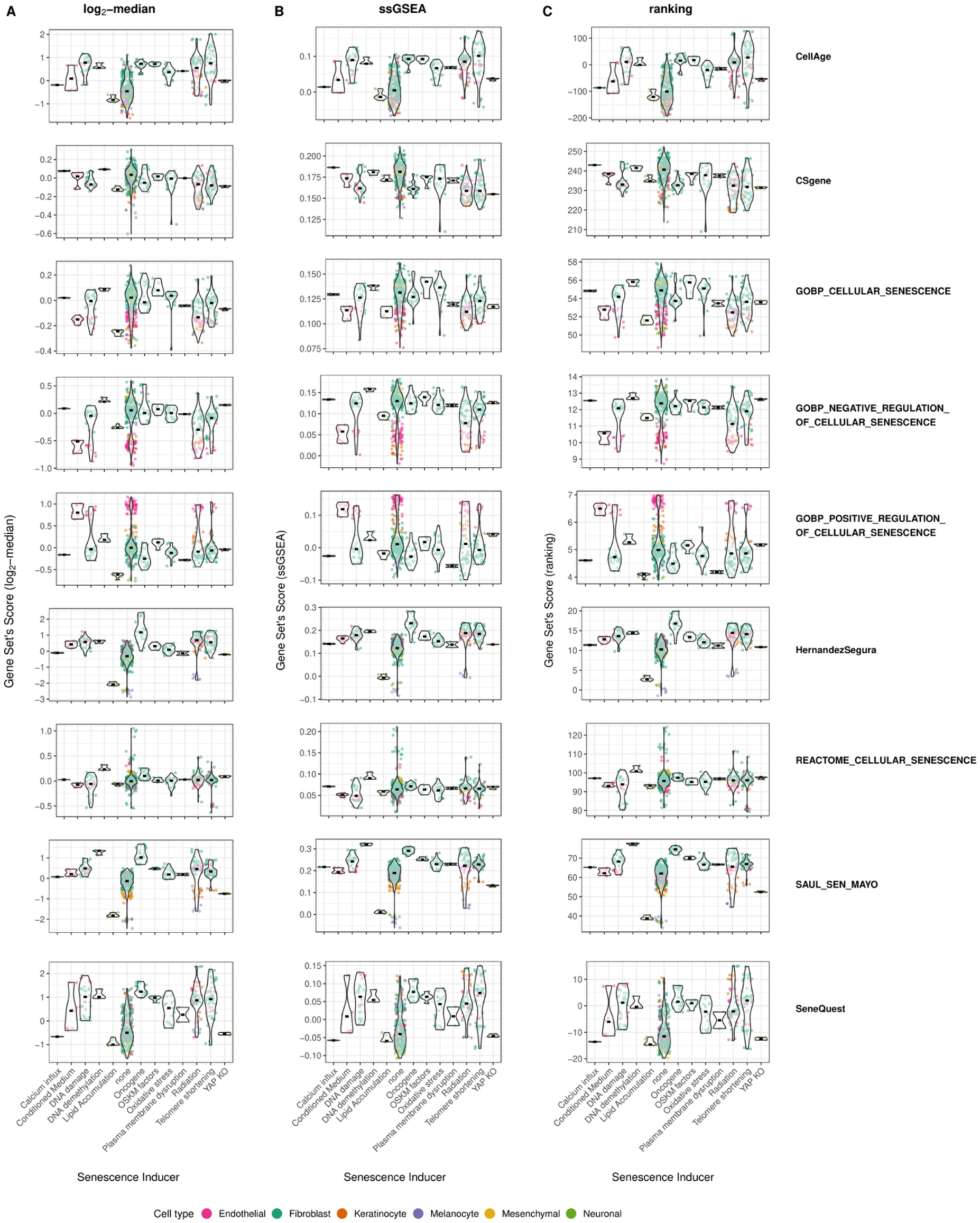
Signature scores across senescence-inducing stressors highlight context specificity. Violin plots showing the distribution of signature scores across different senescence-inducing stressors, calculated using three scoring methods: (A) log_2_-median, (B) ssGSEA, and (C) ranking. Each plot represents one gene signature, with stressor type on the x-axis and score on the y-axis. Data are coloured by cell type. These plots demonstrate that some gene signatures display stressor-specific sensitivity. For example, the HernandezSegura and SAUL_SEN_MAYO signatures are particularly effective in identifying oncogene-induced senescence, and CellAge shows stronger performance in telomere shortening and radiation-induced. In contrast, some signatures show weaker or inconsistent patterns across stressors, highlighting potential limitations in their robustness.

**Supplementary Figure 5.**
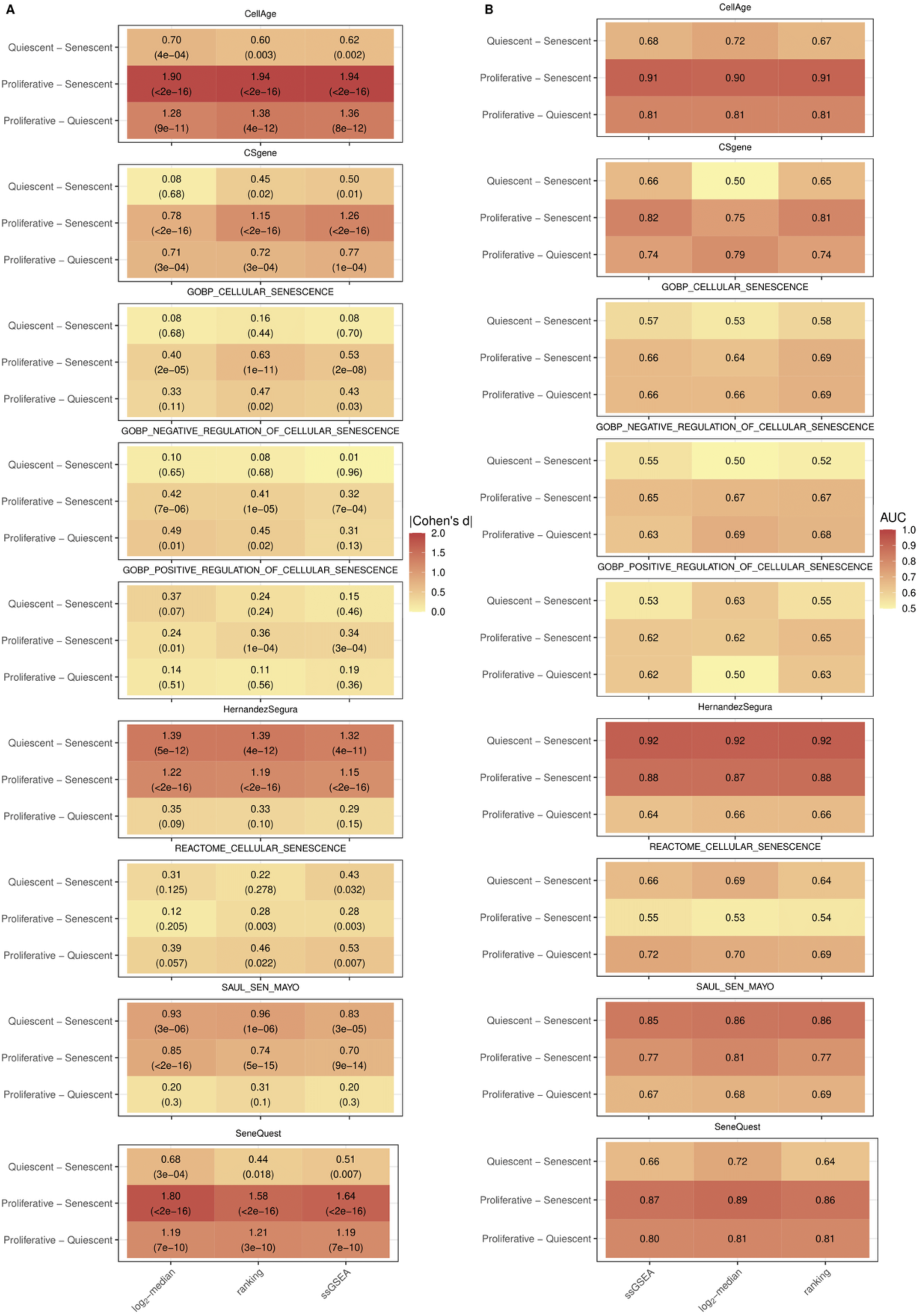
Classification metrics of score-based phenotype quantification methods across gene signatures. (A) Heatmap of absolute effect sizes (Cohen’s d) for each gene signature, with rows representing pairwise contrasts between senescence states and columns representing the scoring methods. A robust signature is expected to show high and consistent effect sizes across methods, particularly for the senescence vs. proliferative and senescence vs. quiescence contrasts. In brackets, the statistical significance given by p-values of the pairwise contrasts’ t-tests. (B) Heatmap of AUC values from ROC curve analyses for the same pairwise contrasts and methods, providing a classification-based evaluation of signature performance.

**Supplementary Figure 6.**
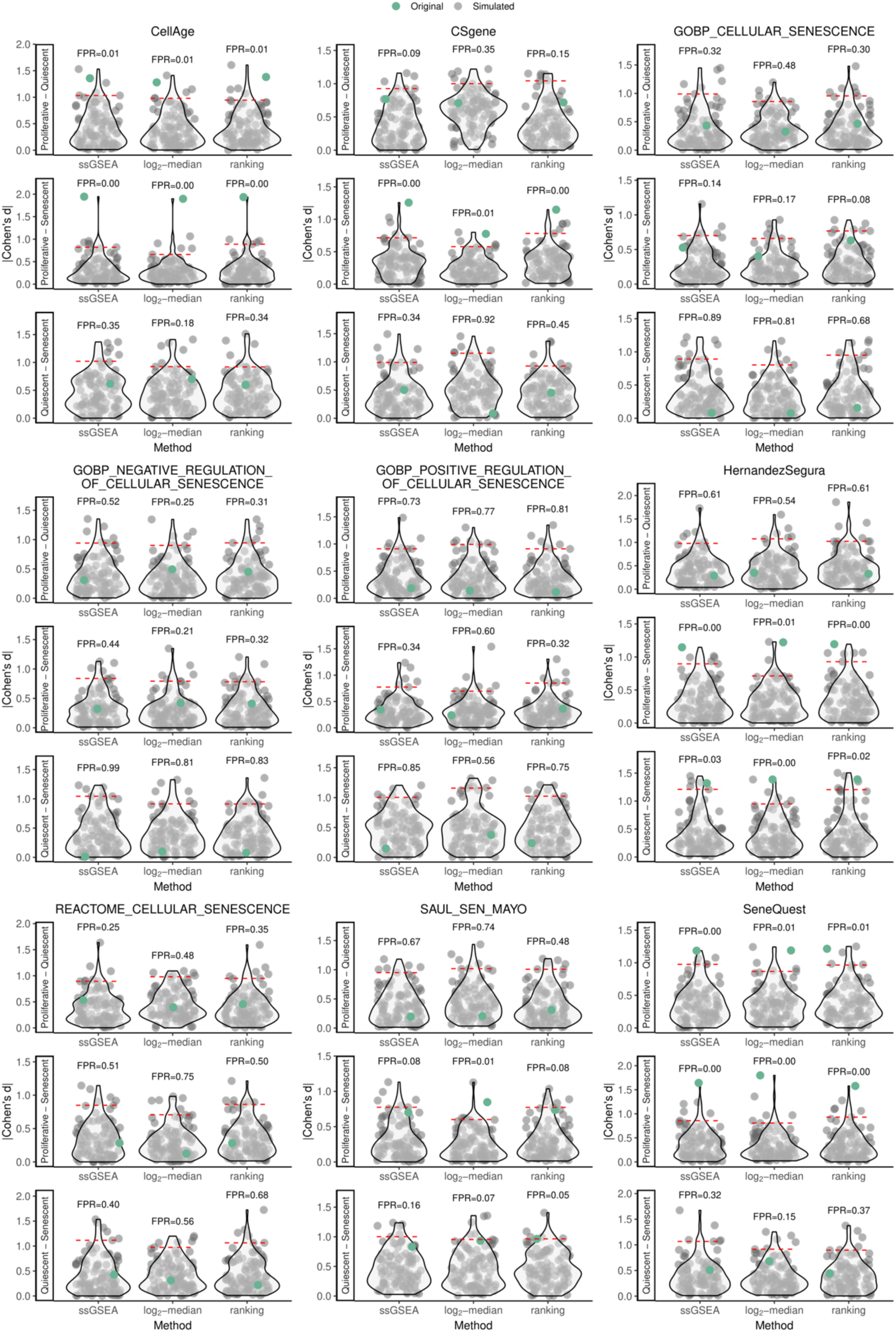
False positive rates (FPRs) across senescence gene signatures using simulated null distributions. The figure is organised into nine groups, with one group representing each gene signature tested. Within each group, three violin plots illustrate the following contrasts: Senescent versus Quiescent, Senescent versus Proliferative, and Proliferative versus Quiescent. For each contrast, the x-axis shows the various scoring methods used. The violin plots show the distribution of absolute Cohen’s d effect sizes from 100 simulated signatures (grey dots) that are matched in size to the real signature, preserving the original signature’s proportion of up- and down-regulated genes, if applicable. Green dots represent the effect size for the actual signature. The dashed red line marks the 95th percentile of the null distribution. False positive rates (FPRs), displayed above each plot, show the proportion of random signatures that outperformed the real signature in each case.

**Supplementary Figure 7.**
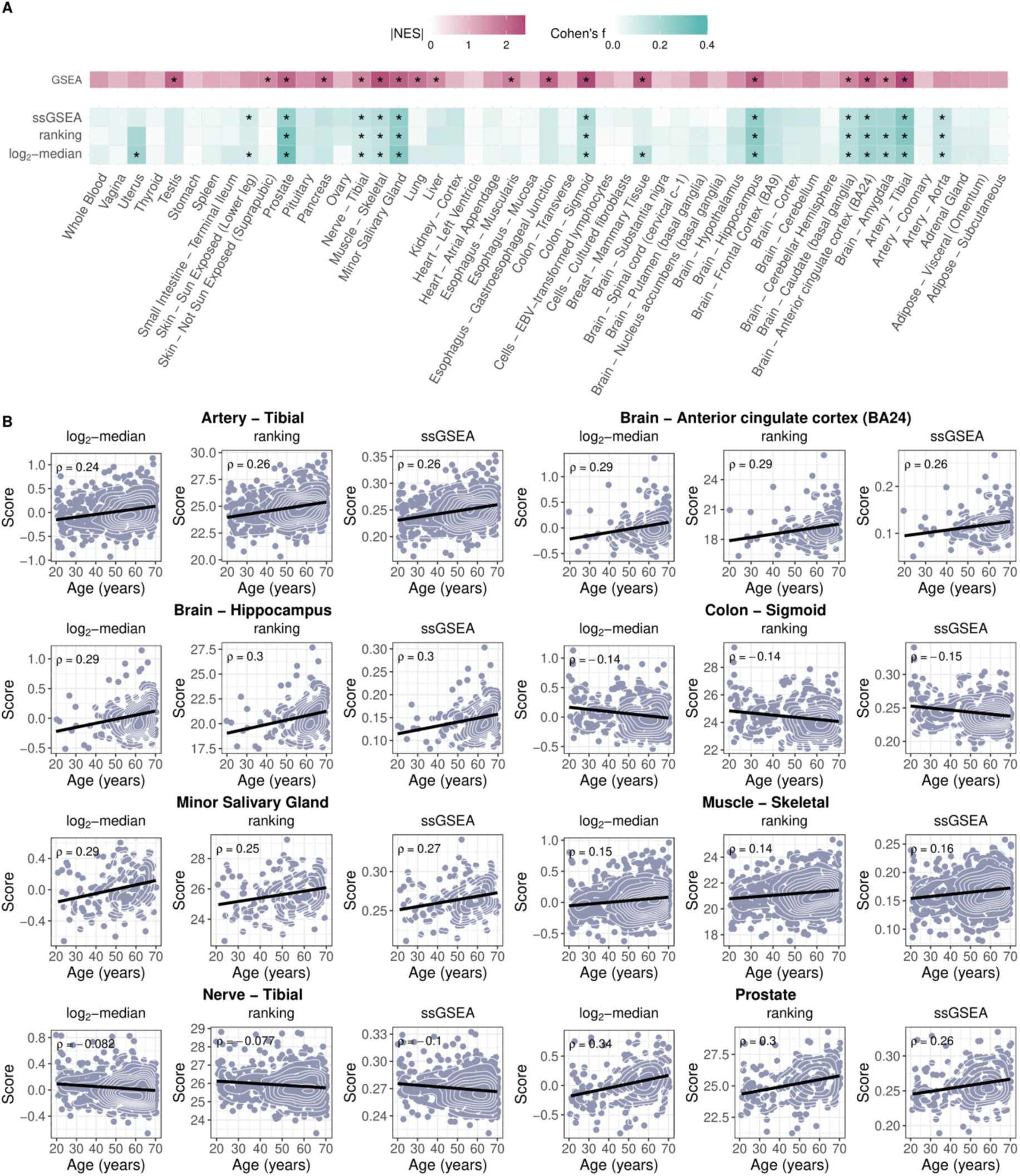
Age-associated senescence scoring and enrichment across GTEx tissues using the SAUL_SEN_MAYO signature. (A) Heatmap showing the association between donors’ age and senescence scores across GTEx tissues, using the SAUL_SEN_MAYO signature. Columns correspond to tissues, and rows represent scoring-based approaches (log2-median, ranking and ssGSEA) using Cohen’s f as a metric of variable association (white to green), and GSEA-based enrichment analysis (NES, white to bordeaux). Asterisks denote significant associations (FDR ≤ 0.05). For the score-based approaches, p-values were obtained from 1,000 random permutations of the age variable, whereas for GSEA, the p-value is derived from the method’s internal enrichment testing procedure. (B) Scatter plots showing score vs age for the eight tissues with significant associations across all four methods in (A). For each tissue, plots for the log2-median, ranking and ssGSEA methods are presented. The age in years (x-axis) is jittered for donor privacy. Black linear regression line and Spearman’s correlation coefficient ρ are shown. Density contour lines in white aid visualisation of dot distribution. Only genes detectable across all 49 tissues were retained for this analysis; the impact of this filtering strategy is shown in Supplementary Figure 11.

**Supplementary Figure 8.**
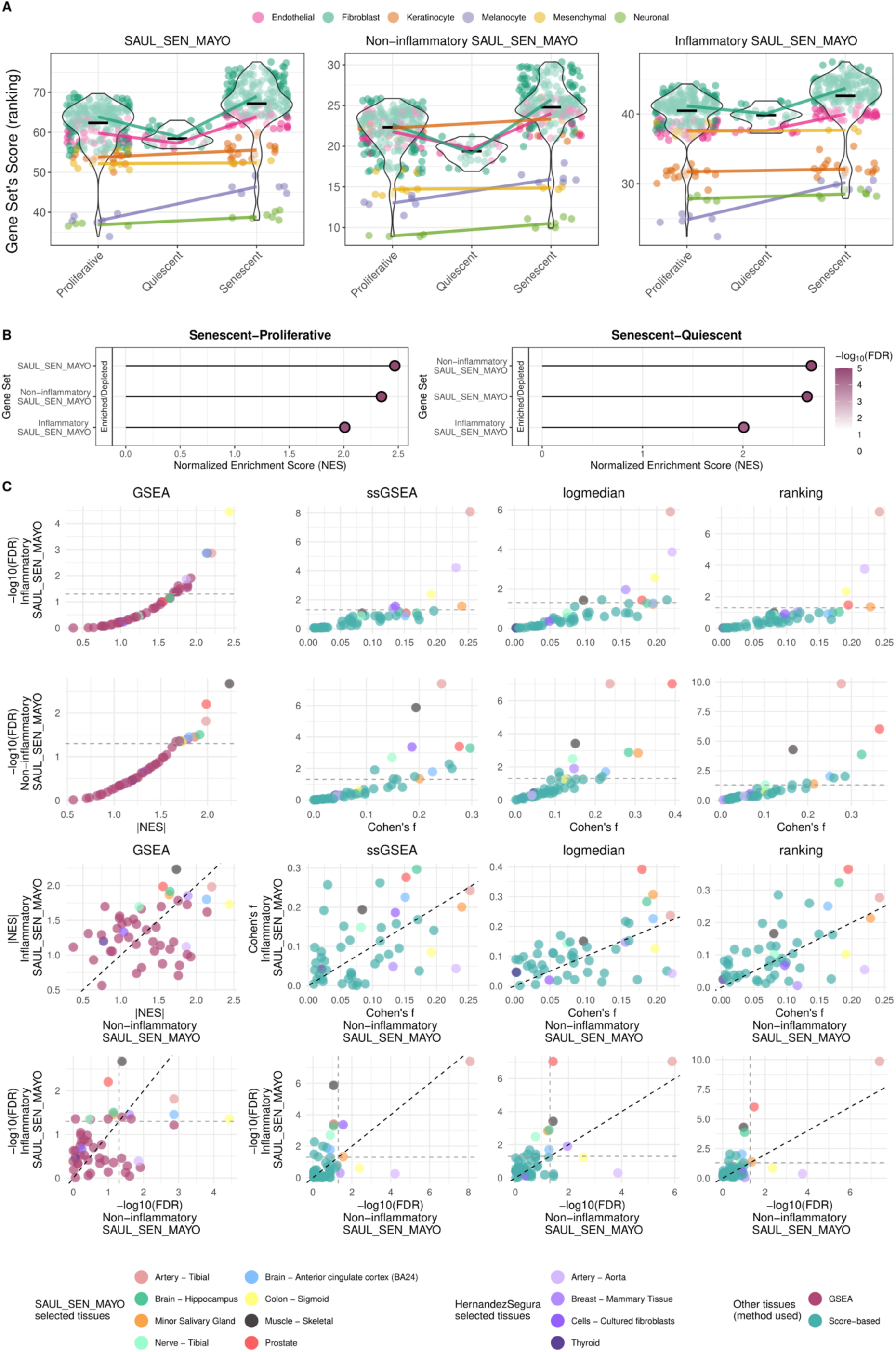
Comparison of inflammatory and non-inflammatory components of the SAUL_SEN_MAYO gene set. (A) Violin plots showing the distribution of scores calculated using the ranking method for three gene sets: the original SAUL_SEN_MAYO signature, the non-inflammatory subset (genes not overlapping with inflammatory Hallmark pathways), and the inflammatory subset (genes overlapping with these pathways). The subsets were defined by intersecting the original gene set with Hallmark gene sets of inflammatory response, TNF-α signalling via NF-κB, interferon gamma and alpha responses, and IL6/JAK/STAT3 signalling. Each plot displays score distributions across sample types (proliferative, quiescent, senescent), coloured by cell type. Median scores per condition and cell type are connected by coloured lines. (B) Gene Set Enrichment Analysis results shown as lollipop plots summarising NES for the three gene sets in the in vitro senescence datasets. The colour gradient represents FDRs, with more intense colours indicating greater statistical significance. (C) Volcano plots (first two rows, showing statistical significance vs. effect size) and correlation plots of effect size (third row, |NES| for GSEA, Cohen’s f for score-based) and statistical significance (fourth row) comparing the inflammatory SAUL_SEN_MAYO versus the non-inflammatory SAUL_SEN_MAYO gene subsets, for the four methods used in markeR (columns: GSEA, ssGSEA, log2-median, ranking). Coloured dots indicate tissues of interest: tissues scored for SAUL_SEN_MAYO highlighted in the main text, tissues scored for Hernandez-Segura in purple shades, and remaining tissues coloured by the method used (teal for score-based, bordeaux for GSEA). For comparisons involving p-values, a dashed grey line marks the 0.05 significance threshold. Black dashed line indicates y=x.

**Supplementary Figure 9.**
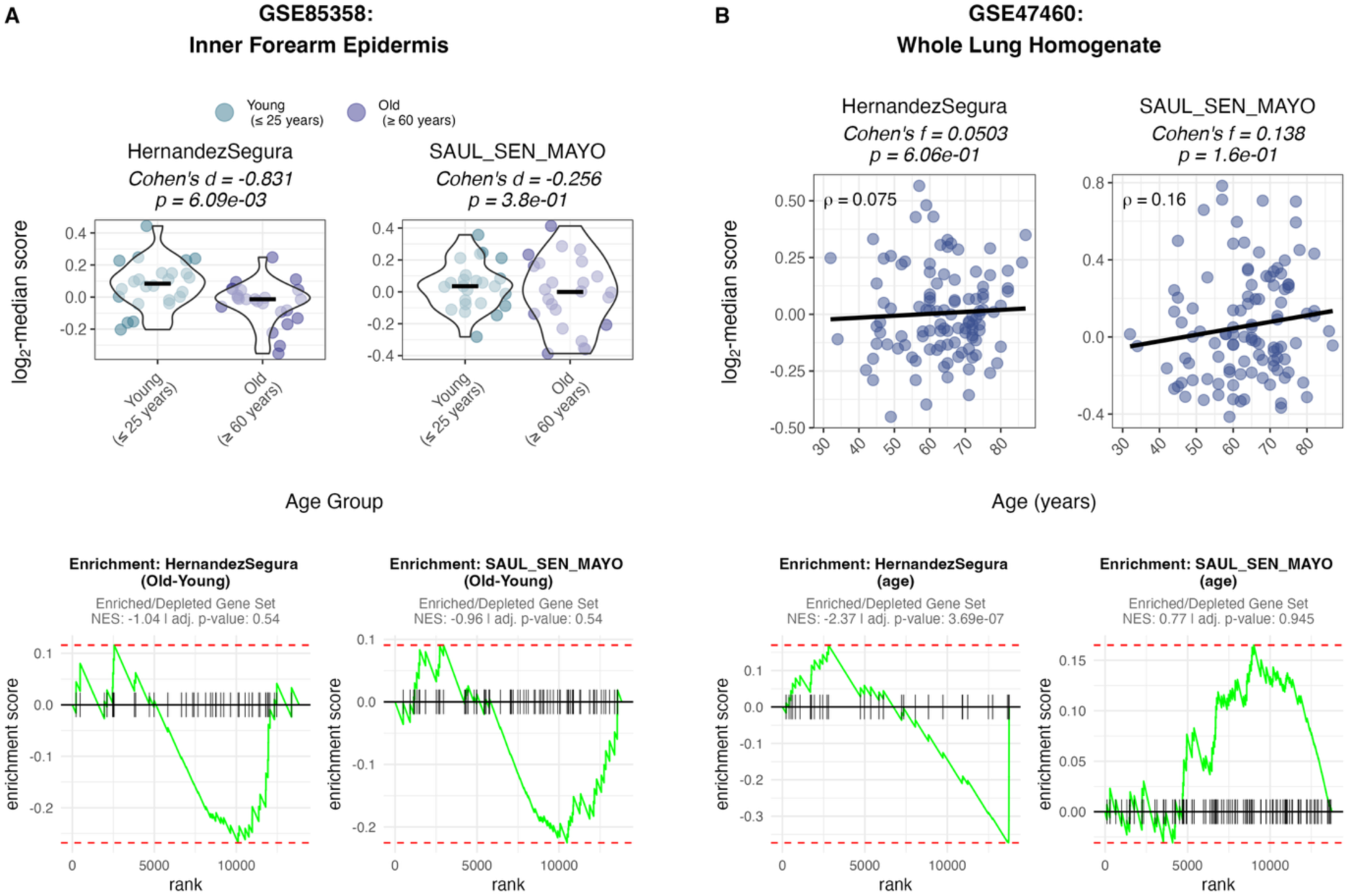
Senescence gene set scores in independent human tissue gene expression datasets across age. **(A)** Inner forearm epidermis (GSE85358; microarray; n = 48; Young ≤ 25 years (n=24), Old ≥ 60 years (n=24)). **(B)** Whole lung homogenate (GSE47460; microarray; n = 108 samples; age range 30–87 years). Microarray data were processed analogously to RNA-seq datasets; low-expression probes were filtered using a minimum total signal threshold of 10^3.2^ (skin) and 10^3.7^ (lung), and probe-level signals were collapsed to gene-level estimates by mean aggregation followed by mapping to HGNC symbols. Scores were computed using the log_2_-median method for two senescence gene sets, HernandezSegura and SAUL_SEN_MAYO. Top row: gene set log_2_-median scores stratified by age group (A) and against age as a continuous variable (B), with a black linear regression line and Spearman’s correlation coefficient ⍴ being shown for the latter. Bottom row: respective GSEA plots, with genes ranked by the t-statistic for differential expression between Old and Young (A) or linearly modelled across age as a continuous variable (B).

**Supplementary Figure 10.**
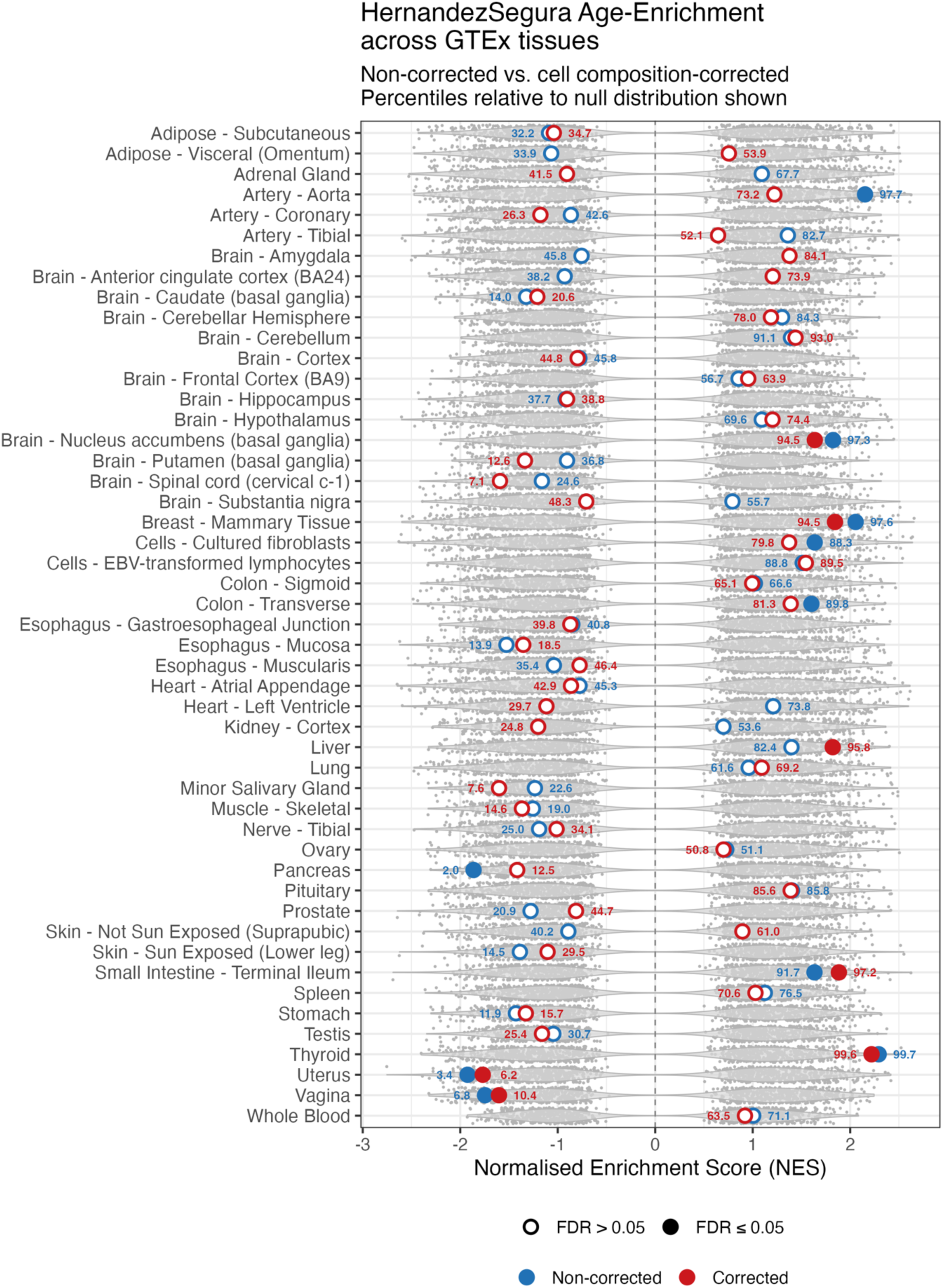
**Effect of cell type composition correction on age-associated HernandezSegura enrichment across GTEx tissues**. For each tissue, NES from the original age-only and the cell type composition-corrected models, the latter incorporating the six most variable xCell2-estimated cell type scores with age per tissue as covariates, are represented by blue and red dots, respectively. The grey violin plots show the tissue-specific null distribution of NES values obtained from 1000 age-label permutations. The number adjacent to each coloured dot indicates its percentile rank within this null distribution (signed, from most negative to most positive). The null distributions reveal a characteristic void of NES values near zero, meaning that visually apparent shifts in NES between models do not necessarily reflect meaningful biological changes; the percentile rank, which contextualises each observed NES within its tissue-specific null distribution, shows that most apparently large shifts remain near the centre of the distribution and are therefore not distinguishable from chance. Tissues are ordered alphabetically. Filled circles indicate FDR ≤ 0.05; open circles indicate FDR > 0.05, following Benjamini-Hochberg correction applied separately per model across all 49 tissues. All genes available in the original processed data for each tissue independently were considered for this analysis.

**Supplementary Figure 11.**
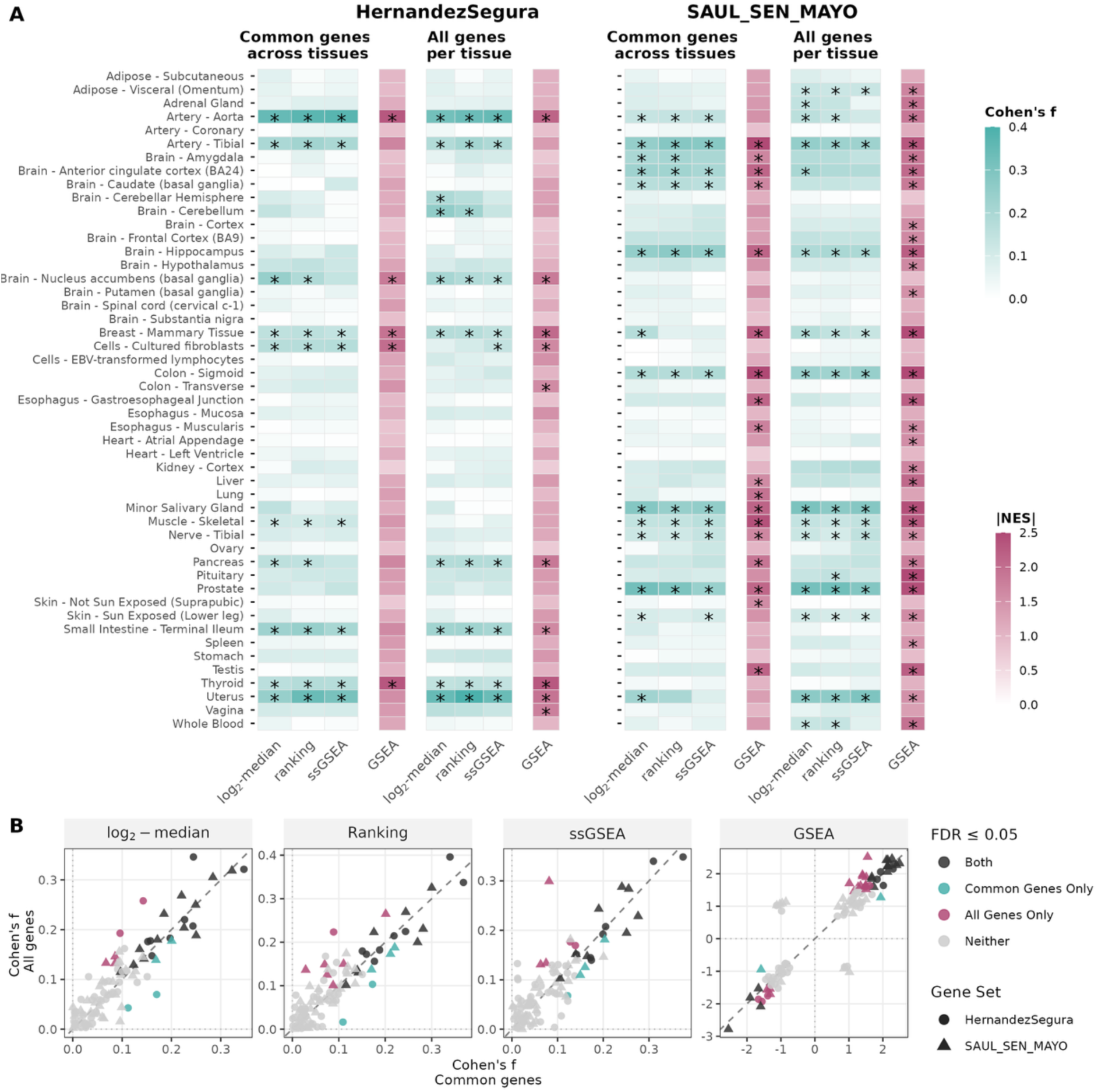
Impact of gene filtering strategy on age-associated senescence signature scoring and enrichment across GTEx tissues. (A) Heatmaps summarising the association between donor age and senescence signature scores (Cohen’s f, teal) and GSEA enrichment (|NES|, pink) across 49 GTEx tissues, for the HernandezSegura (left) and SAUL_SEN_MAYO (right) gene sets. Results are shown under two gene filtering strategies: Common genes across tissues, in which only genes detected across all 49 tissues were retained (as used in the main analysis, *i.e.,* Figure 7 and Supplementary Figures 7 and 8), and All genes per tissue, in which all genes detectable in each tissue were used independently (as required for cell type deconvolution analyses, *i.e.,* Supplementary Figure 9). While retaining only genes common across tissues enables direct cross-tissue comparisons, it may exclude tissue-specific genes that contribute to local senescence signals; the impact of this trade-off is assessed here. Asterisks denote significant associations (FDR ≤ 0.05). (B) Scatter plots directly comparing effect sizes obtained under the two filtering strategies for each scoring method (log₂-median, Ranking, ssGSEA) and GSEA, with each dot representing one tissue-gene set combination. Dots are coloured by significance (FDR ≤ 0.05) concordance between strategies: both significant (dark), common genes only (teal), all genes only (pink), or neither (grey). Dot shape distinguishes the two gene sets. The dashed diagonal is the identity line (y = x). Overall, the two filtering strategies yield highly concordant results across methods and signatures, with only minor fluctuations in significance.

**Supplementary Figure 12.**
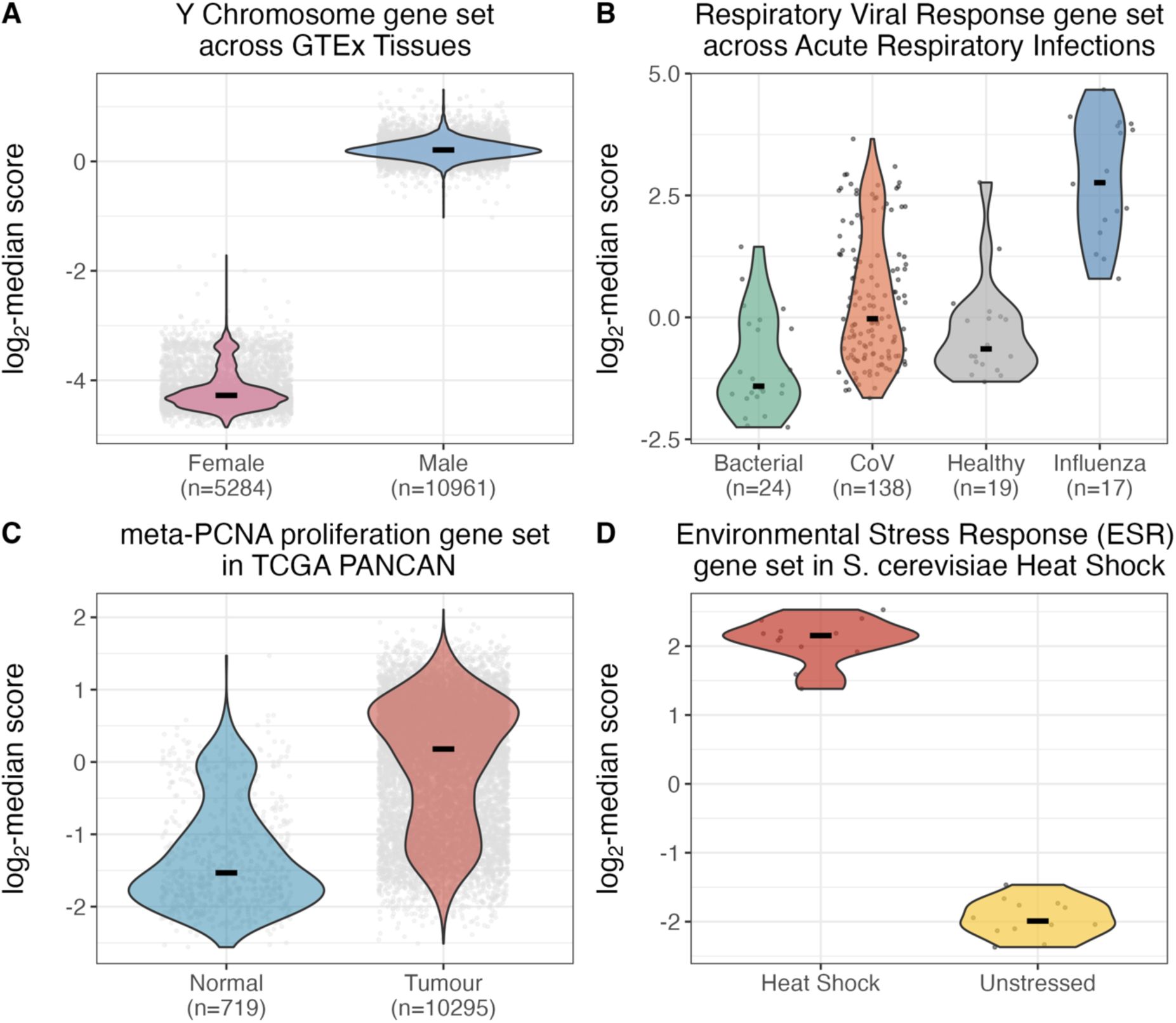
log_2_-median scores for diverse gene sets in relevant biological contexts and organisms. **(A)** A Y chromosome gene set was derived programmatically using the Ensembl BioMart database (accessed via the biomaRt R package (1,2)), by retrieving all protein-coding genes annotated to chromosome Y and excluding those with a homologue on chromosome X. GTEx v8 data (3), already aligned, filtered, normalised and batch-corrected (4) were used directly as input to markeR for scoring across 49 tissues, stratified by donor sex. **(B)** The Respiratory Viral Response gene set (5) comprises peripheral blood genes derived from controlled human viral challenge studies, in which healthy adult volunteers were experimentally inoculated with rhinovirus (HRV), respiratory syncytial virus (RSV), or influenza A, and gene expression was profiled by microarray at peak symptomatic timepoints versus pre-inoculation baseline. Critically, no coronavirus was included in the derivation or validation cohorts. As SARS-CoV-2, for example, is known to actively suppress type I interferon signalling (6), the primary biological process captured by this signature, reduced performance in CoV relative to influenza is mechanistically expected. The signature also distinguishes viral from bacterial infection. Raw counts from GSE161731 (7), already aligned by the original authors, were processed as described in the Methods for the senescence datasets, retaining genes with a total read sum above 10^4.2^ across all samples prior to normalisation. **(C)** The meta-PCNA proliferation gene set (8) comprises genes consistently co-expressed with PCNA across normal human tissues and was shown to underlie most prognostic transcriptional signals in breast cancer. As uncontrolled proliferation is a hallmark of malignant transformation, tumour samples are expected to score systematically higher than matched normal tissue. The signature was applied here to The Cancer Genome Atlas (TCGA) PANCAN data (9), comprising RNA-seq profiles across 33 cancer types, accessed as pre-processed expression matrices through the UCSC Xena Browser (10), to compare scores between primary tumour and matched normal tissue samples. **(D)** The Environmental Stress Response (ESR) gene set (11) comprises two transcriptionally coordinated gene groups defined across 142 microarray experiments in Saccharomyces cerevisiae: an induced component (iESR, ∼300 genes) upregulated upon stress, and a repressed component (rESR, ∼600 genes) downregulated upon stress. The gene set was retrieved from the neat R package (12), and the combined bidirectional set was applied to RNA-seq data from GSE135430 (13) in which S. cerevisiae BY4741 cells grown at 30°C were subjected to 20 minutes of heat shock at 37°C. Pre-processed expression data provided by the original authors were used directly, retaining genes with a total read sum above 10^3.4^ across all samples prior to normalisation. As heat shock is one of the canonical ESR-inducing conditions, a strong increase in the bidirectional ESR score in heat-shocked relative to unstressed cells is expected.

## Notes

https://github.com/DiseaseTranscriptomicsLab/markeR

### Competing Interest Statement

The authors have declared no competing interest.

### Summary of Updates

Clarified GTEx analysis strategy, specifying gene filtering across tissues, and added new supplementary figure comparing alternative filtering approaches; Incorporated cell type deconvolution in GTEx analyses and included estimated proportions as covariates to address confounding effects; Added systematic comparison with existing gene set scoring methods (e.g., GSVA, UCell) and clarified the positioning and added value of markeR; Expanded Discussion to address methodological limitations, broader applicability, and adaptation to single-cell data; Implemented a lightweight Python interface (via rpy2) enabling access to markeR functionality without requiring R coding; Improved clarity of methodological descriptions, figure annotations, and overall presentation throughout the manuscript.

